# Bayesian Estimation of Population Size Changes by Sampling Tajima’s Trees

**DOI:** 10.1101/605352

**Authors:** Julia A. Palacios, Amandine Véber, Lorenzo Cappello, Zhangyuan Wang, John Wakeley, Sohini Ramachandran

## Abstract

The large state space of gene genealogies is a major hurdle for inference methods based on Kingman’s coalescent. Here, we present a new Bayesian approach for inferring past population sizes which relies on a lower resolution coalescent process we refer to as “Tajima’s coalescent”. Tajima’s coalescent has a drastically smaller state space, and hence it is a computationally more efficient model, than the standard Kingman coalescent. We provide a new algorithm for efficient and exact likelihood calculations for data without recombination, which exploits a directed acyclic graph and a correspondingly tailored Markov Chain Monte Carlo method. We compare the performance of our Bayesian Estimation of population size changes by Sampling Tajima’s Trees (BESTT) with a popular implementation of coalescent-based inference in BEAST using simulated data and human data. We empirically demonstrate that BESTT can accurately infer effective population sizes, and it further provides an efficient alternative to the Kingman’s coalescent. The algorithms described here are implemented in the `R` package `phylodyn`, which is available for download at https://github.com/JuliaPalacios/phylodyn.

## 1 Introduction

Modeling gene genealogies from an alignment of sequences — timed and rooted bifurcating trees reflecting the ancestral relationships among sampled sequences — is a key step in coalescent-based inference of evolutionary parameters such as effective population sizes. In the neutral coalescent model without recombination, observed sequence variation is produced by a stochastic process of mutation acting along the branches of the gene genealogy (Kingman, 1982a; Watterson, 1975), which is modeled as a realization of the coalescent point process at a neutral non-recombining locus. In the coalescent point process, the rate of coalescence (the merging of two lineages into a common ancestor at some time in the past) is a function that varies with time, and it is inversely proportional to the effective population size at time *t, N*(*t*) (Kingman, 1982b; Slatkin and Hudson, 1991; Donnelly and Tavaré, 1995). Our goal is to infer (*N*(*t*))_*t*≥0_ which we will refer to as the “effective population size trajectory”.

Multiple methods have been developed to infer (*N*(*t*))_*t*≥0_ using the standard coalescent model with or without recombination. Some of these methods infer (*N*(*t*))_*t*≥0_ from summary statistics such as the sample frequency spectrum (SFS) (Terhorst et al., 2017; Bhaskar et al., 2015); however, the SFS is not a sufficient statistic for inferring (*N*(*t*))_*t*≥0_ (Sainudiin et al., 2011). Other methods have been proposed that directly use molecular sequence alignments at a completely linked locus, i.e. without recombination (Griffiths and Tavaré, 1996; Kuhner and Smith, 2007; Minin et al., 2008; Li and Durbin, 2011; Drummond et al., 2012; Palacios and Minin, 2013; Gill et al., 2013). Our approach is of this type. Still other methods account for recombination across larger genomic segments (Li and Durbin, 2011; Sheehan et al., 2013; Schiffels and Durbin, 2014; Palacios et al., 2015). In spite of their variety, all these methods must contend with two major challenges: (*i*) choosing a prior distribution or functional form for (*N*(*t*))_*t*≥0_, and (*ii*) integrating over the large hidden state space of genealogies. For example, several previous approaches have assumed exponential growth (Griffiths and Tavaré, 1996; Kuhner et al., 1998; Kuhner and Smith, 2007), in which case the estimation of (*N*(*t*))_*t*≥0_ is reduced to the estimation of one or two parameters. In general, the functional form of (*N*(*t*))_*t*≥0_ is unknown and needs to be inferred. A commonly used naive nonparametric prior on (*N*(*t*))_*t*≥0_ is a piecewise linear or constant function defined on time intervals of constant or varying sizes (Heled and Drummond, 2008; Sheehan et al., 2013; Schiffels and Durbin, 2014). The specification of change points in such time-discretized effective population size trajectories is inherently difficult because it can lead to runaway behavior or large uncertainty in 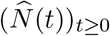. These difficulties can be avoided by the use of Gaussian-process priors in a Bayesian nonparametric framework, allowing accurate and precise estimation (Palacios and Minin, 2013; Gill et al., 2013; Lan et al., 2015; Palacios et al., 2015). More precisely, the autocorrelation modeled with the Gaussian process avoids runaway behavior and large uncertainty in 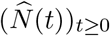.

The second challenge for coalescent-based inference of (*N*(*t*))_*t*≥0_ is the integration over the hidden state space of genealogies for large sample sizes. Given molecular sequence data **Y** at a single non-recombining locus and a mutation model with vector of parameters ***μ***, current methods rely on calculating the marginal likelihood function Pr(**Y**|(*N*(*t*))_*t*≥0_, ***μ***) by integrating over all possible coalescence and mutation events. Under the infinite-sites mutation model without intra-locus recombination (Watterson, 1975), this integration requires a computationally expensive importance sampling technique or Markov Chain Monte Carlo (MCMC) techniques (Griffiths and Tavaré, 1994a; Stephens and Donnelly, 2000; Hobolth et al., 2008; Wu, 2010; Gronau et al., 2011). Moreover, a maximum likelihood estimate of (*N*(*t*))_*t*≥0_ cannot be explicitly obtained; instead, it is obtained by exploring a grid of parameter values (Tavaré, 2004). For finite-sites mutation models, current methods approximate the marginal likelihood function by integrating over all possible genealogies via MCMC methods (Equation (1); Kuhner (2006); Drummond et al. (2012)). In both cases, the marginal likelihood may be expressed as

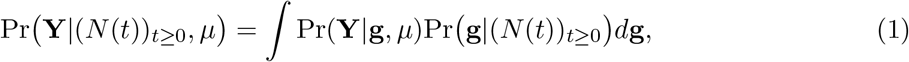

in which Pr(·) is used to denote both the probability of discrete variables and the density of continuous variables. The integral above involves an (*n* − 1)-dimensional integral over *n* − 1 coalescent times and a sum over all possible tree topologies with *n* leaves. Therefore, these methods require a very large number of MCMC samples, and exploration of the posterior space of genealogies continues to be an active area of research (Kuhner et al., 1998; Rannala and Yang, 2003; Drummond et al., 2012; Whidden and Matsen, 2015; Aberer et al., 2016).

Current methods rely on the Kingman *n*-coalescent process to model the sample’s ancestry. However, the state space of genealogical trees grows superexponentially with the number of samples, making inference computationally challenging for large sample sizes. In this study, we develop a Bayesian nonparametric model that relies on Tajima’s coalescent, a lower resolution coalescent process with a drastically smaller state space than that of Kingman’s coalescent. In particular, we approximate the posterior distribution Pr((*N*(*t*))_*t*≥0_, **g**^*T*^, ***τ*** | **Y**, *μ*), where **g**^*T*^ corresponds to the Tajima’s genealogy of the sample (see Figure 1A and Section 2.4), (log *N*(*t*))_*t*≥0_ has Gaussian process prior with precision hyperparameter τ that controls the degree of regularity, and mutations occur according to the infinite-sites model of Watterson (1975). This results in a new efficient method for inferring (*N*(*t*))_*t*≥0_ called **B**ayesian **E**stimation by **S**ampling **T**ajima’s **T**rees (BESTT), with a drastic reduction in the state space of genealogies. We show using simulated data that BESTT can accurately infer effective population size trajectories and that it provides a more efficient alternative than Kingman’s coalescent models.

**Figure 1:**
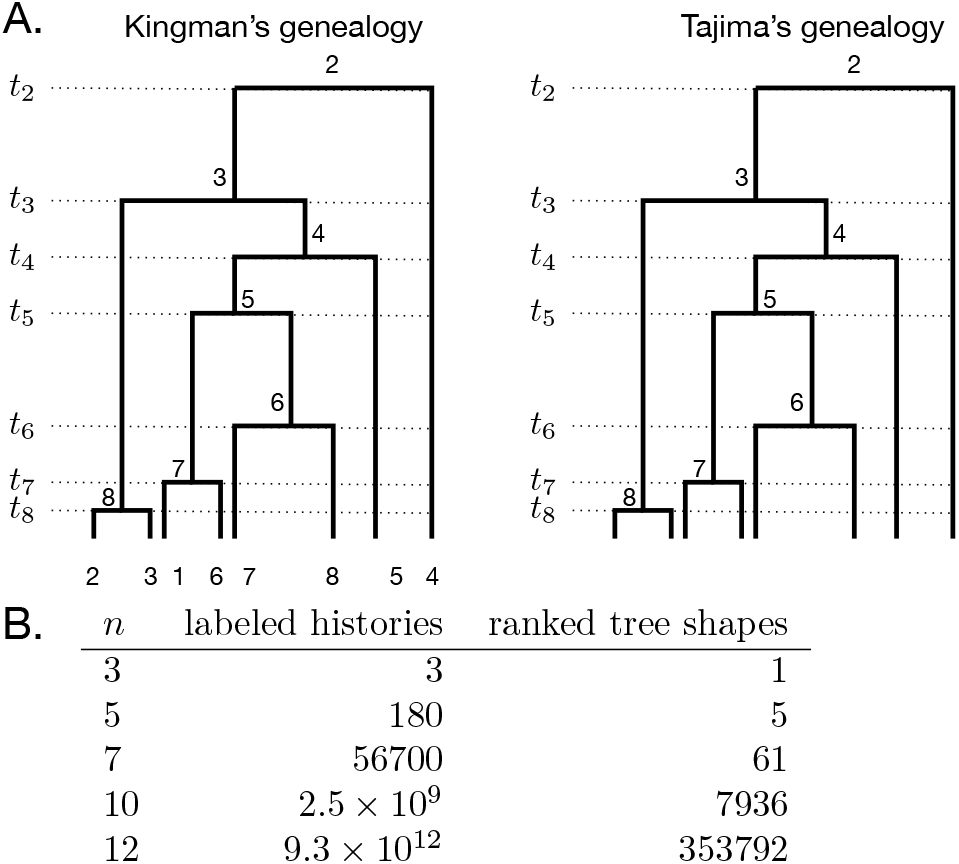
For a sample of size *n*, the number of Tajima’s genealogies is su-perexponentially fewer compared to the number of Kingman’s genealogies. **A**: A Kingman’s genealogy and a Tajima’s genealogy for *n* = 8. A Kingman’s genealogy (left) comprises a vector of coalescent times and the labeled topology; the number of possible labeled topologies for a sample of size *n* is *n*!(*n* − 1)!/2^*n*−1^. A Tajima’s genealogy (right) comprises a vector of coales-cent times and a ranked tree shape. In both cases, coalescent events are ranked from 2 at time *t*_2_ to *n* at time *t_n_*. Coalescent times are measured from the present (time 0) back into the past. **B**. The numbers of labeled topologies and ranked tree shapes (formulas provided in section 2.4) for different values of the sample size, *n*.

Next, we start with an overview of BESTT, detail our representation of molecular sequence data and define the Tajima coalescent process. We then introduce a new augmented representation of sequence data as a directed acyclic graph (DAG). This representation allows us to both calculate the conditional likelihood under the Tajima coalescent model, and to sample tree topologies compatible with the observed data. We then provide an algorithm for likelihood calculations and develop an MCMC approach to efficiently explore the state space of unknown parameters. Finally, we compare our method to other methods implemented in BEAST (Drummond et al., 2012) and estimate the effective population size trajectory from human mtDNA data. We close with a discussion of possible extensions and limitations of the proposed model and implementation.

## 2 Methods/Theory

### 2.1 Overview of BESTT

Our objective in the implementation of BESTT is to estimate the posterior distribution of model parameters by replacing Kingman’s genealogy with Tajima’s genealogy **g**^*T*^. A Tajima’s genealogy does not include labels at the tips (Figure 1): we do not order individuals in the sample but label only the lineages that are ancestral to at least two individuals (that is, we only label the internal nodes of the genealogy). Replacing Kingman’s genealogy by Tajima’s genealogy in our posterior distribution exponentially reduces the size of the state space of genealogies (Figure 1B). In order to compute Pr(**Y**|**g**^*T*^, *μ*), the conditional likelihood of the data conditioned on a Tajima’s genealogy, we assume the infinite sites model of mutations and leverage a directed acyclic graph (DAG) representation of sequence data and genealogical information. Note that the overall likelihood, Eq. (1), will differ only by a combinatorial factor from the corresponding likelihood under the Kingman coalescent. Our DAG represents the data with a gene tree (Griffiths and Tavaré, 1994a), constructed via a modified version of the perfect phylogeny algorithm of Gusfield (1991). This provides an economical representation of the uncertainty and conditional independences induced by the model and the observed data.

Under the infinite-sites mutation model, there is a one-to-one correspondence between observed sequence data and the gene tree of the data (Gusfield, 1991) (Sections 2.2–2.3). We further augment the gene tree representation with the allocation of the number of observed mutations along the Tajima’s genealogy to generate a DAG (Section 2.5). The conditional likelihood Pr(**Y** | **g**^*T*^, *μ*) is then calculated via a recursive algorithm that exploits the auxiliary variables defined in the DAG nodes, marginalizing over all possible mutation allocations (Section 2.6). We approximate the joint posterior distribution Pr((*N*(*t*))_*t*≥0_, **g**^*T*^, ***τ*** | **Y**, *μ*) via an MCMC algorithm using Hamiltonian Monte Carlo for sampling the continuous parameters of the model and a novel Metropolis-Hastings algorithm for sampling the discrete tree space.

### 2.2 Summarizing sequence data Y as haplotypes and mutation groups

Let the data consist of *n* fully linked haploid sequences or alignments of nucleotides at *s* segregating sites sampled from *n* individuals at time *t* = 0 (the present). Note that any labels we afix to the individuals are arbitrary in the sense that they will not enter into the calculation of the likelihood. We further assume the infinite sites mutation model of Watterson (1975) with mutation parameter *μ* and known ancestral states for each of the sites. Then we can encode the data into a binary matrix **Y** of *n* rows and *s* columns with elements *y_i,j_* ∈ {0,1}, where 0 indicates the ancestral allele.

In order to calculate the Tajima’s conditional likelihood Pr(**Y** | **g**^*T*^, *μ*), we first record each haplotype’s frequency and group repeated columns to form *mutation groups*; a mutation group corresponds to a shared set of mutations in a subset of the sampled individuals. We record the cardinality of each mutation group (*i.e*., the number of columns that show each mutation group). In Figure 2A, there are two columns labeled “b”, corresponding to two segregating sites which have the exact same pattern of allelic states across the sample. Further, two individuals carry the derived allele of mutation group “b”, so in this case the frequency of haplotype 7 and the cardinality of mutation group “b” are both equal to 2. Likewise, haplotype 4 has frequency 1 and carries 5 mutations that are split into mutation groups “a”, “f” and “g” (the latter is not shown in Figure 2A, but appears in Figure 2B) of respective cardinalities 1, 3 and 1. We denote the number of haplotypes in the sample as *h*, the number of mutation groups as *m*, and the representation of **Y** as haplotypes and mutation groups as **Y**_*h*×*m*_.

**Figure 2:**
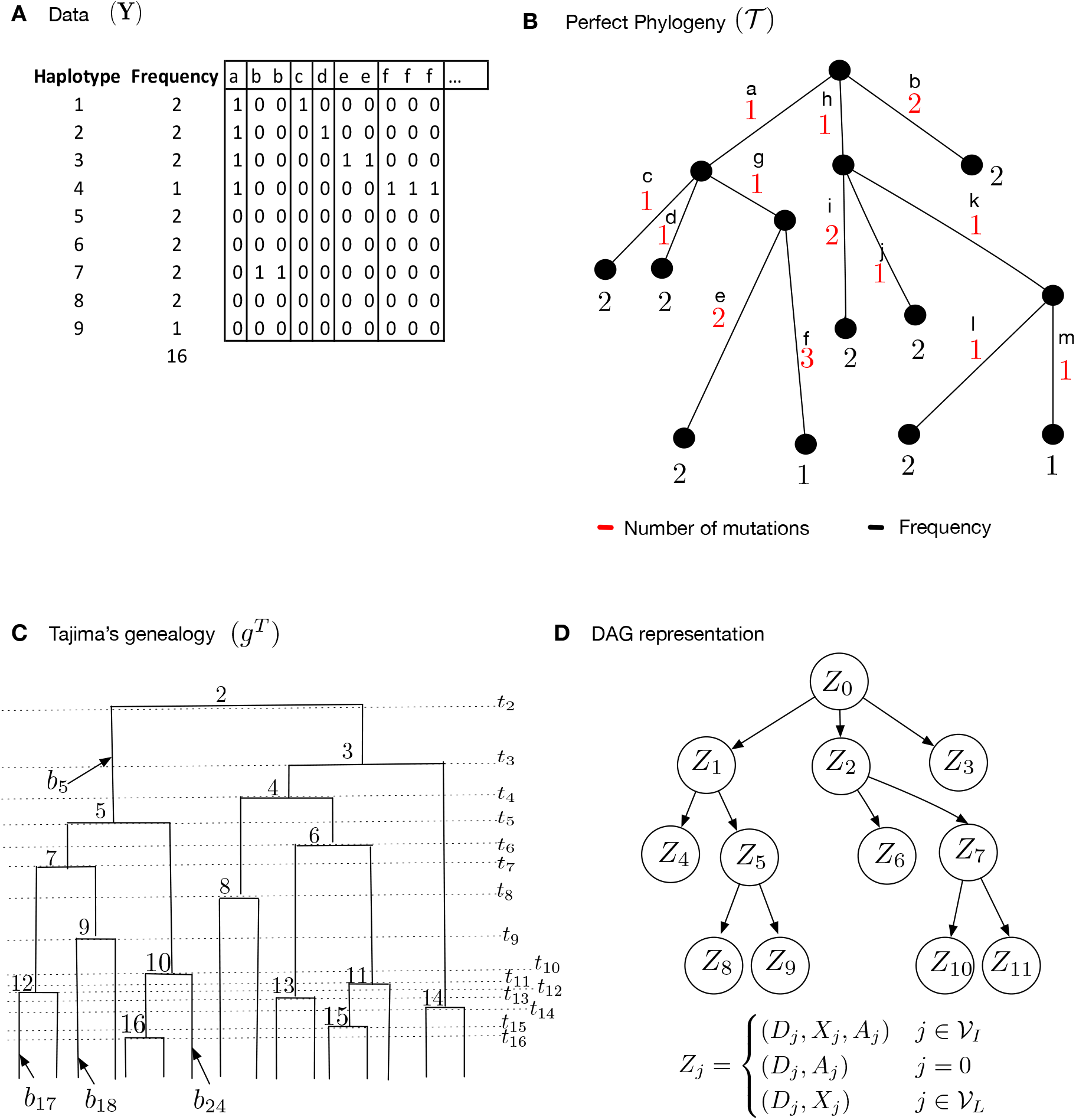
Data structures employed by our method, BESTT, for calculating the conditional likelihood of the data. **A.** Compressed data representation **Y**_*h*×*m*_ of *n* = 16 sequences and *s* = 18 (columns, only the first 10 of which are shown), comprised of 9 haplotypes and 13 mutation groups. Rows correspond to haplotypes and each polymorphic site is labeled by its mutation group {*a, b, c*,…, *m*}. **B.** Gene tree representation of the data in panel A. Red numbers indicate the cardinality of each mutation group (number of columns with the same label in panel A). Black letters indicate the mutation group (column labels in panel A), and black numbers indicate the frequency of the corresponding haplotype. **C.** A Tajima’s genealogy compatible with the gene tree in panel B. Internal nodes are labeled according to order of coalescent events from the root to the tips. Coalescent event *i* happens at time *t_i_* and branches are labeled *b_i_* (see section 2.5 for details). **D.** A Directed Acyclic Graph (DAG) representation of the gene tree in panel B together with allocation of mutation groups along the branches of the Tajima’s genealogy in panel C. 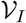 denotes the set of internal nodes and 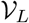 the set of leaf nodes. A detailed description of the DAG is given in section 2.5.

### 2.3 Representing Y_*h*×*m*_ as a gene tree

**Y**_*h*×*m*_ (Figure 2A) can alternatively be represented as a gene tree or perfect phylogeny (Gusfield, 1991; Griffiths and Tavaré, 1994b). This representation relies on our assumption of the infinite sites mutation model in which, if a site mutates once in a given lineage, all descendants of that lineage also have the mutation *and no other individuals carry that mutation*. The gene tree is a graphical representation of the haplotypes (as tips) arranged by their patterns of shared mutations. The haplotype data summarized in Figure 2A corresponds to the gene tree given in Figure 2B. Details of the correspondence between haplotype data and gene tree are listed below, and an additional example is given in Figure 13 (Appendix E).

A *gene tree* for a matrix **Y**_*h*×*m*_ of *h* haplotypes and *m* mutation groups is a rooted tree 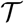 with *h* leaves and at least *m* edges, such that (Figure 2B):

1. Each row of **Y**_*h*×*m*_ corresponds to exactly one leaf of 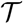. The black numbers at leaf nodes in Figure 2B are the haplotype frequencies.
2. Each mutation group of **Y**_*h*×*m*_ is represented by exactly one edge of 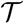, which is labeled accordingly (letters in Figures 2A and 2B). The red numbers along edges in Figure 2B give the cardinality of each mutation group (*i.e*. the number of segregating sites in this group; see Figure 2A). Some external edges (edges subtending leaves) may not be labeled, indicating that they do not carry additional mutations to their parent edge. This happens when the other edges emanating from its parent node necessarily correspond to other mutation groups.
3. Edges are placed in the gene tree in such a way that each path from the root to a leaf fully describes a haplotype. Edges corresponding to shared mutations between several haplotypes are closest to the root. For example, in Figure 2B, haplotype 4 corresponds to the leaf at which one arrives starting from the root and going along edges a, g and f; in contrast, haplotype 7 corresponds to the leaf at which one arrives going from the root along edge b. Thus, the labels and the numbers associated with the edges along the unique path from the root to a leaf exactly specify a row of **Y**_*h*×*m*_.

Dan Gusfield’s perfect phylogeny algorithm (Gusfield, 1991) transforms the sequence data **Y**_*h*×*m*_ into a gene tree and this transformation is one-to-one. We note that the perfect phylogeny 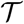 or gene tree is not the same as the genealogy g. While a genealogy is a bifurcating tree of individuals of the sample, the gene tree is a multifurcating tree of haplotypes.

### 2.4 Tajima’s genealogies

Our method of computing the probability of the recoded data, **Y**_*h*×*m*_, uses ranked tree shapes rather than fully labeled histories. We refer to these ranked tree shapes as Tajima’s genealogies but note they have also been called *unlabeled rooted trees* (Griffiths and Tavaré, 1995) and *evolutionary relationships* (Tajima, 1983). In Tajima’s genealogies, only the internal nodes are labeled and they are labeled by their order in time. Tajima’s genealogies encode the minimum information needed to compute the probability of data, **Y**_*h*×*m*_ which consists of nested sets of mutations, without any approximations. In Figure 1A for example, it matters only that mutation group “e” occurs on a subgroup of the individuals who carry a mutation group “a” and that this is different than the subgroups carrying “c”, “d” and “f”. No other labels matter because individuals are exchangeable in the population model we assume.

This represents a dramatic coarsening of tree space compared to the classical leaf-labeled binary trees of Kingman’s coalescent. The number of possible ranked tree shapes for a sample of size *n* corresponds to the *n*-th term of the sequence A000111 of Euler zig-zag numbers (Disanto and Wiehe, 2013) whereas the number of labeled binary tree topologies is *n*!(*n* − 1)!/2^*n*−1^. As can be seen from Figure 1B, this provides a much more efficient way to integrate over the key hidden variable, the unknown gene genealogy of the sample, when computing likelihoods.

We model this hidden variable using the *vintaged and sized coalescent* (Sainudiin et al., 2015) which corresponds exactly to this coarsening of Kingman’s coalescent. As can be seen in Figure 1A, we assign vintages/labels 2 through *n* starting at the root of the tree and moving toward the present, so that the node created by the final splitting event, which is also the first coalescence event looking back in the ancestry of the sample, is labeled *n*. We write *t_k_* for the time of node *k*, measured from the present back into the past. We set *t*_*n*+1_: = 0 to be the present time. Then during the interval [*t*_*k*+1_, *t_k_*) the sample has exactly *k* extant ancestors, for *k* ∈ {2,…, *n*}.

The coarsening of the tree topology does not change the law of the times between two coalescence events. Thus, conditional on the effective population size trajectory (*N*(*t*))_*t*≥0_ and the time *t*_*k*+1_ at which the number of ancestors to the sample decreases to *k*, the distribution of the time during which the sample has *k* ancestors is given by

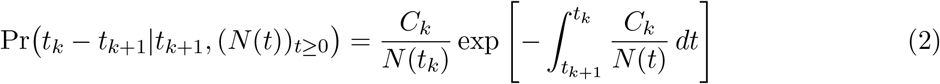

(Slatkin and Hudson, 1991), where 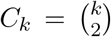. Writing the density at **t** = (*t*_2_, *t*_3_,…, *t_n_*) of the vector of coalescence times as a product of conditional densities, we obtain

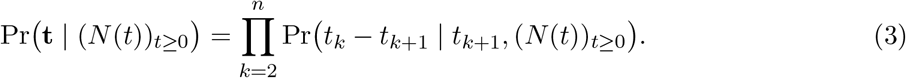

We use a lower-triangular matrix denoted **F** to represent Tajima’s genealogies; see Appendix A. The probability of a ranked tree shape was derived independently in Sainudiin et al. (2015) and Palacios et al. (2015). Specifically, for every ranked tree shape **F** with *n* leaves,

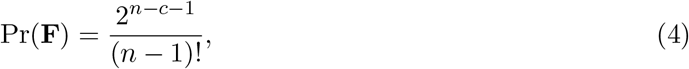

where *c* is the number of cherries in **F** (*i.e*., nodes subtending two leaves; *c* = 3 in Figure 10A). Note that this probability is independent of the effective population size trajectory since the choice of the pair of lineages that coalesce during an event is independent of (*N*(*t*))_*t*≥0_ (recall that in Kingman’s coalescent, the coalescing pair is chosen uniformly at random among all possible pairs). Since the distribution of Tajima’s genealogies **g**^*T*^ = (**F, t**) conditional on (*N*(*t*))_*t*≥0_ can be factored as the product of the probability of the ranked tree shape **F** and the coalescent times density, we arrive at

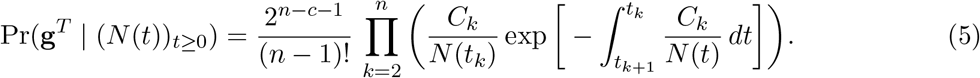

### 2.5 An augmented data representation using directed acyclic graphs

A key component of BESTT is the calculation of the conditional likelihood Pr(**Y**|**g**^*T*^, *μ*). We compute the conditional likelihood recursively over a directed acyclic graph (DAG) **D.** Our DAG exploits the gene tree representation 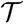 of the data (Figure 2B), incorporates the branch length information of the Tajima’s genealogy **g**^*T*^(Figure 2C) and facilitates the recursive allocation of mutations to the branches of **g**^*T*^. Here we detail the construction of the DAG.

We construct the DAG using three pieces of information: the observed gene tree 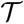, a given Tajima’s genealogy **g**^*T*^ and a latent “allocation” of mutations along the branches of the Tajima’s genealogy (Figure 3). An allocation refers to a possible mapping (compatible with the data) of the observed numbers of mutations (red numbers in Figure 2B) to branches in the Tajima’s genealogy. Figure 3A shows one possible mapping for the Tajima’s genealogy in Figure 2C; usually this mapping is not unique. Our construction of **D** enables an efficient recursive consideration of all possible allocations of mutations along **g**^*T*^ when computing the conditional likelihood Pr(**Y** | **g**^*T*^, *μ*).

**Figure 3:**
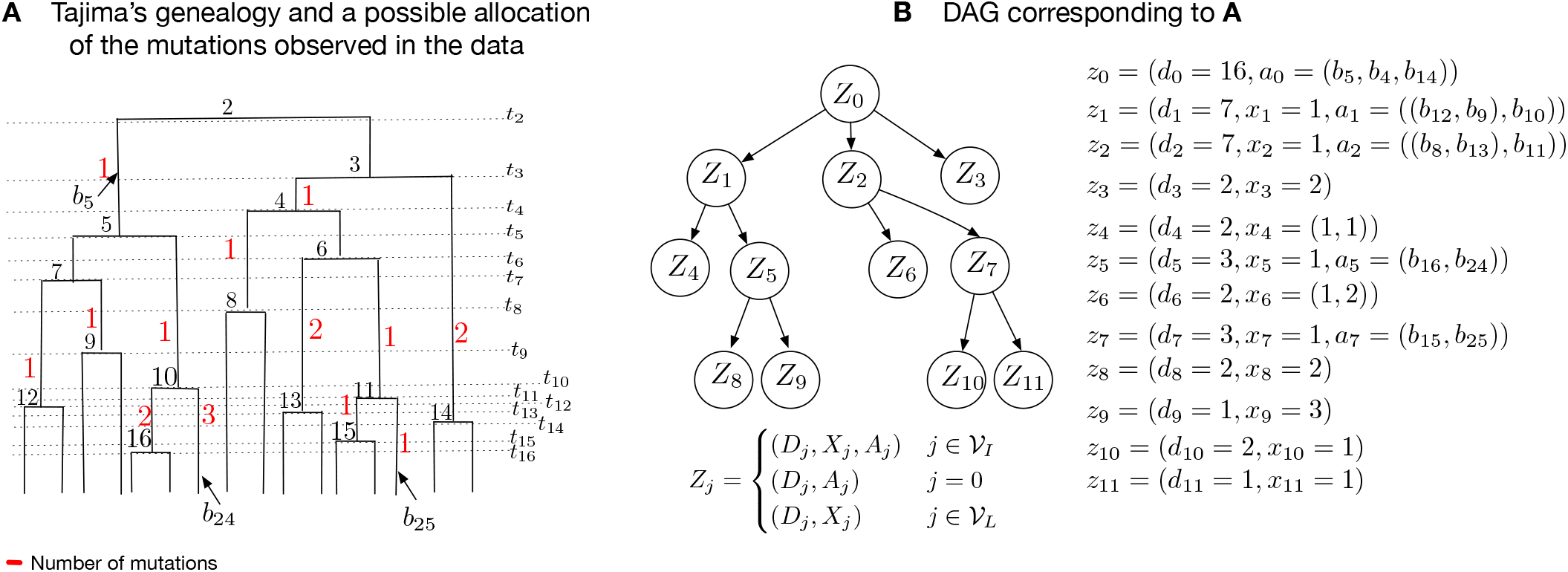
DAG Construction. **A.** A Tajima’s genealogy from Figure 2C with added allocation of mutations shown in red. **B.** The corresponding augmented DAG with allocation of mutations. At the root *Z*_0_, there are no mutations by convention. Node *Z*_0_ has 16 descendants across 3 subtrees of 7, 7 and 2 descendants, corresponding to nodes *Z*_1_, *Z*_2_, *Z*_3_. These three subtrees subtend from *b*_5_, *b*_4_ and *b*_14_, respectively, in **g**^*T*^ (Figure 3A). Node *Z*_1_ corresponds to the tree subtending from *b*_5_ of size 7 with *X*_1_ = 1 mutation along *b*_5_ and subtends three subtrees from (*b*_12_, *b*_9_) and *b*_10_. Subtrees subtending from (*b*_12_, *b*_9_) are grouped together in leaf node *Z*_4_ because they both have 2 descendants and have the same parent node. When leaf nodes represent more than one trees, such as *Z*_4_ in Figure 4B, the random variable *X_j_* is the vector *X_j_* = (*X*_*j*,1_, *X*_*j*,2_,…, *X_j,s_j__*) that denotes the number of mutations along the branches that subtends from the tree node *j* that have *D_j_* descendants, and *s_j_* is the number of edges subtending from *Z_j_*.

#### Constructing the DAG D

The graph structure of our DAG **D** = {**Z**, *E*} (Figure 2D) with nodes **Z** and edges *E* is constructed from a gene tree 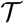. The number of internal nodes in the DAG **D** is the same as the number of internal nodes in 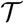. However, sister leaf nodes in 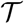 with the same number of descendants are grouped together in **D** and leaf nodes descending from edges with no mutations are treated as singletons grouped together in **D**. For example, the leaves in Figure 2B subtending from edges *i* and *j* are grouped into *Z*_6_ in Figure 2D, as they both have haplotype frequency 2. However, the leaves subtending from the e and f edges are not grouped (and correspond to *Z*_8_ and *Z*_9_ in the DAG Figure 2D) since they have respective haplotype frequencies 2 and 1. We label the root node of **D** as *Z*_0_ and increase the index *i* of each node *Z_i_* from top to bottom, moving left to right. For *i* < *j*, we assign a directed edge *E_i,j_* if the node in 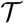 corresponding to *Z_i_* is connected to the node in 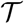 corresponding to *Z_j_*. The index set of internal nodes in **D** is denoted by 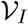 and the index set of leaf nodes is denoted by 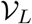.

#### Information carried by the nodes in D

Each node in **D** represents a vector, *Z_j_*, which includes number of descendants, number of mutations and latent allocation of mutations. Although the number of descendants and number of mutations are part of the observed data, the allocation of mutations can be seen as a random variable, for ease of exposition, we use capital letters to denote all three types of information. We define the vector *Z_j_* as follows:

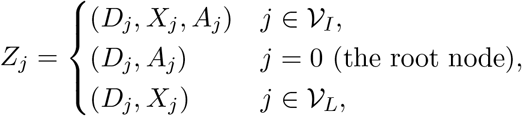

where *D_j_* denotes the number of descendants of (*i.e*., of sampled sequences subtended by) node *Z_j_, X_j_* denotes the number of mutations separating *Z_j_* from its parent node, and *A_j_* denotes the allocation of mutations along **g**^*T*^ (described in detail below). The number of descendants *D_j_* is thus the number of individuals/sequences descending from node *Z_j_* (this information is part of 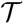). For internal nodes, *X_j_* records the cardinality of a mutation group, represented as a red number along the edge *E_i,j_* of 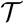 in Figure 2B, where *i* is the index of the parent node of *Z_j_*. Leaf nodes in **D** may correspond to more than one leaf nodes in 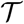, namely any sister nodes with the same number of descendants. In this case, *X_j_* is a vector with the cardinalities of the corresponding mutation groups (see for example node *Z*_6_ in Figure 3B). In order to keep the DAG construction simple, we only allow groupings of leaf nodes and not of internal nodes with identical descendants carrying identical numbers of mutations. We note that, in principle, it would be possible to compress the number of internal nodes of the DAG by exploiting all the symmetries observed in the data.

#### Allocation of mutation groups along g^*T*^

The latent allocation variables {*A_j_*} determine a possible correspondence between subtrees in **g**^*T*^ and nodes in **D**: in particular, *A_j_* indicates the branches in **g**^*T*^ that subtend the subtrees corresponding to nodes {*Z_k_*} if {*Z_k_*} are child nodes of *Z_j_*.

Allocations of mutations to branches are usually not unique and computation of the conditional likelihood Pr(**Y** | **g**^*T*^, *μ*) requires summing over all possible allocations. In Figure 3A we show one such possible allocation of the mutation groups of the gene tree in Figure 2B along the Tajima’s genealogy in Figure 2C. For example, mutation group “a” in Figure 2B with cardinality 1 (number in red) is a mutation observed in 7 individuals (sum of black numbers of leaves descending from edge marked a). This same mutation group, “a”, is shown as a red number 1 in Figure 3A allocated to branch *b*_5_. If *Z_j_* is an internal node, the number of mutations *X_j_* is denoted as a vector of length 1. If *Z_j_* is a leaf node, *X_j_* can be a vector of length greater than 1. Details on notation for allocations can be found in Appendix B.

### 2.6 Computing the conditional likelihood

Under the infinite-sites mutation model, mutations are superimposed independently on the branches of **g**^*T*^ as a Poisson process with rate *μ*. In order to compute 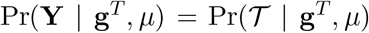 we marginalize over the latent allocation information in the directed acyclic graph **D**; that is, we sum over all possible mappings of mutations in 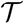 to branches in **g**^*T*^ as follows:

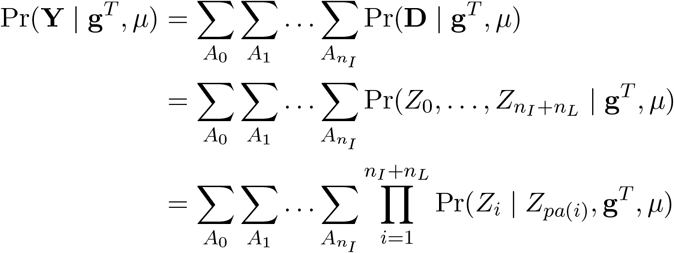

where 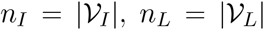, *pa*(*i*) denotes the index of the parent of node *i* in **D** and we set *P*(*Z*_0_ | **g**^*T*^, *μ*) = 1 because it is assumed that there are no mutations above the root node and the length of the root branch *l*_2_ = 0. Writing 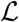 for the tree length of **g**^*T*^ (*i.e*., the sum of the lengths of all branches of **g**^*T*^) and factoring out a global factor 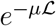 (due to the Poisson distribution of mutations across the genealogy) from each of the above products over *i* ∈ {1,…,*n_I_* + *n_L_*}, we have

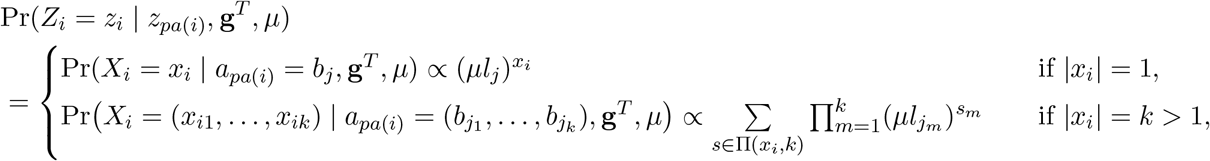

where Π(*x_i_, k*) is the set of all permutations of *x_i_* = {*x*_*i*1_,…, *x_ik_* } divided into *m_i_* groups of different sizes. The number of different permutations of the *k* values of *x_i_* divided into *m_i_* groups of sizes *k*_1_,…, *k_m_i__* is

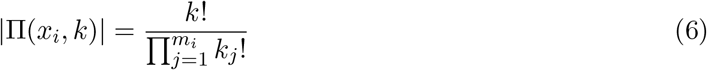

For example, assume that *x_i_* = {2,2,2, 0, 3, 3} and *a*_*pa*(*i*)_ = (*b*_3_, *b*_4_, *b*_5_, *b*_6_, *b*_7_, *b*_8_) with branch lengths {*l*_3_, *l*_4_, *l*_5_, *l*_6_, *l*_7_, *l*_8_}. In this case, *k*_1_ = 3 because there will be 3 branches with 2 mutations, *k*_2_ = 1 because there will be 1 branch with 0 mutations and *k*_3_ = 2 because there will be 2 branches with 3 mutations. The number of permutations of *k* = 6 mutations groups divided into *m_i_* = 3 groups with cardinalities 2, 0, 3 of sizes 3, 1, 2 is 6!/(3!1!2!) = 60.

The conditional likelihood Pr(**Y** | **g**^*T*^, *μ*) is calculated via a backtracking algorithm (Appendix C). The algorithm marginalizes the allocations by traversing the DAG from the tips to the root. The pseudocode and an example can be found in the Appendix C.

### 2.7 The case of unknown ancestral states

Up to now, we have assumed that the ancestral state was known at every segregating site. The representation of the data **Y** that we use in this case records the cardinalities of each mutation group and the genealogical relations between these groups, but does not assign labels to the sequences. Hence, in the terminology of Griffiths and Tavaré (1995), our data corresponds to an *unlabeled rooted gene tree*.

When the ancestral types are not known, the data (now denoted **Y**^0^) may be represented as an unlabeled *unrooted* gene tree. By the remark following Equation (1) in Griffiths and Tavaré (1995), if *s* is the number of segregating sites, then there are at most *s* + 1 unlabeled rooted gene trees that correspond to the unrooted gene tree of the observed data 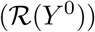. By the law of total probability (see also Equation (10) in Griffiths and Tavaré (1995)), the conditional likelihood of **Y**^0^ can be written as the sum over all compatible unlabeled rooted gene trees *Y*^(*i*)^ of the probability of *Y*^(*i*)^ conditionally on **g**^*T*^. That is:

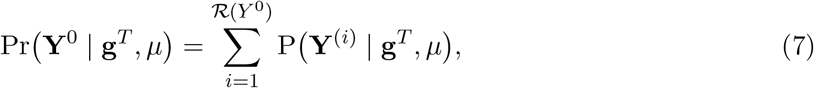

where each of the **Y**^(*i*)^ corresponds to a unique unlabeled rooted gene tree compatible with the unrooted gene tree **Y**^0^ and 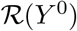 denotes the number of those unlabeled rooted gene trees. In the following sections, we shall assume that the ancestral type at each site is known.

### 2.8 Bayesian inference of the effective population size trajectory

Our posterior distribution of interest is

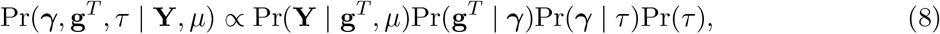

where 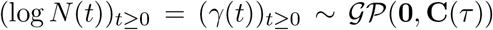 has a Gaussian process prior with mean **0** and covariance function **C**(*τ*) (Rasmussen and Williams, 2006). This specification ensures (*N*(*t*))_*t*≥0_ is non-negative. In our implementation, we assume a regular geometric random walk prior, that is, 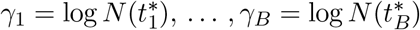 at *B* regularly spaced time points in [0, *T*] with

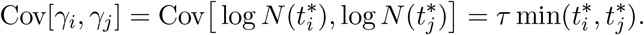

The parameter *τ* is a length scale parameter that controls the degree of regularity of the random walk. We place a Gamma prior with parameters *α* = .01 and *β* = .001 on *τ*, reflecting our lack of prior information in terms of high variance about the smoothness of the logarithm of the effective population size trajectory.

We approximate the posterior distribution of model parameters via a MCMC sampling scheme. Model parameters are sampled in blocks within a random scan Metropolis-within-Gibbs framework. Our algorithm initializes with the corresponding Tajima genealogy of the UPGMA estimated tree implemented in `phangorn` (Schliep, 2011). Given an initial genealogy, our algorithm initializes *N_e_* and *τ* with the method of (Palacios and Minin, 2012) implemented in `phylodyn` (Karcher et al., 2017). We then proceed to generate (1) a sample of the vector of effective population sizes and precision parameter as described in section 2.8.2, (2) a sample of the vector of coalescent times as described in 2.8.3 and 2.8.4 where we modify a single coalescent time, and (3) a sample of ranked tree shape as described in 2.8.1 in each iteration. To summarize the effective population size trajectory, we compute the posterior median and 95% credible intervals pointwise at each grid point in 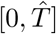, were 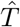 is the maximum time to the most recent common ancestor sampled.

#### 2.8.1 Metropolis-Hastings updates for ranked tree shapes

There is a large literature on local transition proposal distributions for Kingman’s topologies (Kuhner et al., 1998; Rannala and Yang, 2003; Drummond et al., 2012; Whidden and Matsen, 2015; Aberer et al., 2016). In this paper, we adapted the local transition proposal of Markovtsova et al. (2000) to Tajima’s topologies. We briefly describe the scheme below and provide a pseudocode algorithm in Appendix C (Algorithm 3).

Given the current state of the chain {*γ, τ*, **g**^*T*^} = {*γ,τ*, **F**_*n*_, **t**}, we propose a new ranked tree shape **F**\* in two steps: (1) we first sample a coalescent interval *e_k_* = (*t*_*k*+1_, *t_k_*) uniformly at random, where *k* ~ U({3,…, *n*}). Note that we will never select the interval (*t*_3_, *t*_2_) at the top of the tree (see Figure 10A). Given *k*, we focus solely on the coalescent events at times *t_k_* and *t*_*k*−1_. For step (2), there are two possible scenarios. Case A: The lineage created at time *t_k_*, labeled *k*, coalesces at time *t*_*k*−1_ (first row of Figure 4A). Case B: Lineage *k* does not coalesce at time *t*_*k*−1_ (Figure 4B). In Case A, we choose a new pair of lineages at random to coalesce at time *t_k_* from the 3 lineages subtending *k* and *k* − 1 (excluding *k*), and we coalesce the remaining lineage with *k* at *t*_*k*−1_ (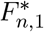 and 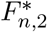 in Figure 4). In Case B, we invert the order of the coalescent events; that is, the two lineages descending from *k* are set to coalesce at time *t*_*k*−1_ and lineages descending from *k* − 1 are set to coalesce at time *t_k_*. (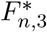 a in Figure 4). Note that the numerical labels 1, 2,3 are included to clarify the picture: lineages subtending both Case A and Case B can be either labeled (if there is a vintage subtending that lineage) or not (if there is a singleton). The transition probability 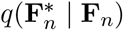 is given by the product of the probabilities of the two steps. The new ranked tree shape 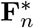 is accepted with probability given by the Metropolis-Hastings ratio defined below:

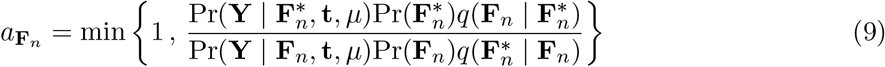

**Figure 4:**
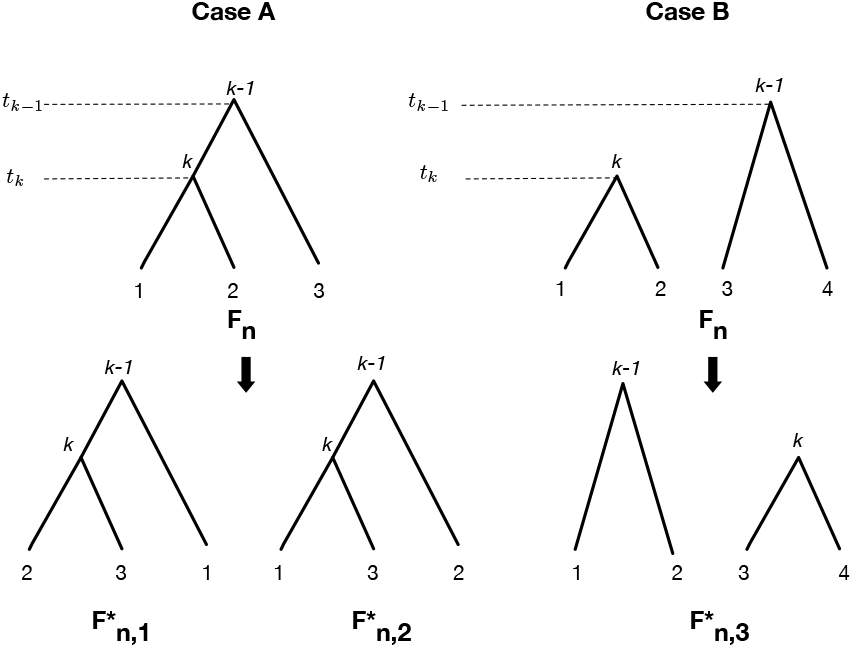
Markov moves for topologies. First row: possible coalescent patterns (Case A or Case B) for a given topology *F_n_*. Second row: possible Markov moves in Case A (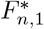 and 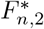) and Case B 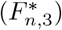. *k* indexes the coalescent interval sampled. Numerical labels at the tips are added for convenience: conditionally on a given *F_n_*, tips can be labeled (vintage) or not (singleton). Figure is adapted from Figures 2, 3 and 4 of Markovtsova et al. (2000).

We note that our proposal can result in the same ranked tree shape. However, we tested alternative proposals that precluded this event and we did not find any notable difference in the overall performance of the MCMC algorithm.

#### 2.8.2 Split Hamiltonian Monte Carlo updates of (*γ, τ*)

To make efficient joint proposals of *γ* and *τ*, we use the Split Hamiltonian Monte Carlo method proposed by Lan et al. (2015). Conditioned on *g^T^*, the target density becomes *π*(*γ, τ*) ∝ Pr(**t** | *γ*)Pr(*γ* | *τ*)Pr(*τ*). This is the same target density implemented in Karcher et al. (2017) for fixed coalescent times **t**.

#### 2.8.3 Hamiltonian Monte Carlo updates of coalescent times

Given the current state {*γ, τ*, **g**^*T*^} = {*γ*, **F**_*n*_, **t**, *τ*}, we propose a new vector of coalescent times with target density *π*(**t**′) ∝ P(**Y** | **F**_*n*_, **t**′, *μ*)P(**t**′ | *γ*) by numerically simulating a Hamilton system with Hamiltonian

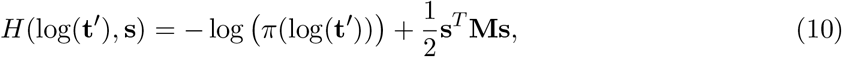

where **s** is the momentum vector assumed to be normally distributed. The system evolves according to:

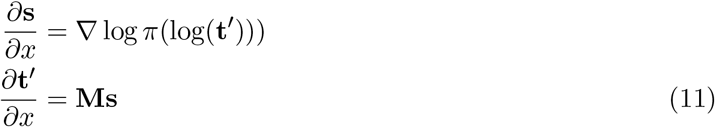

We use the *leapfrog* method (Neal, 2011) with step size *ϵ* and a *p* Poisson with mean 10 distributed number of steps to simulate the dynamics from time *x* = 0 to *x* = *pϵ*. Each leapfrog step of size *ϵ* follows the trajectory:

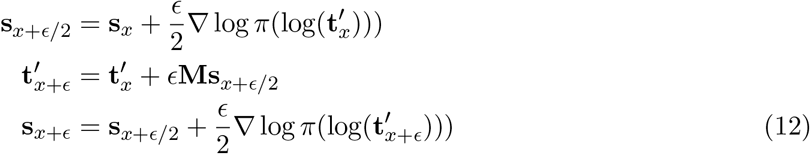

For our implementation, we set the mass matrix **M** = **I**, the identity matrix. We simulate the Hamiltonian dynamics of the logarithm of times to avoid proposals with negative values. Solving the equations of the Hamilton system requires calculating the gradient of the logarithm of the target density with respect to the vector of log coalescent times. The gradient of the log conditional likelihood (score function) is calculated at every marginalization step in the algorithm for the likelihood calculation.

At the beginning of Section 2.8, we described how we assume a regular geometric random walk prior on (*N*(*t*))_*t*≥0_ at *B* regularly spaced time points in [0,*T*]. Ideally, the window size *T* must be at least *t*_2_, the time to the most recent common ancestor (TMRCA). However, *t*_2_ is not known. Our initial values of coalescent times *t* are obtained from the UPGMA implementation in phangorn (Schliep, 2011) with times properly rescaled by the mutation rate, and we set *T* = *t*_2_. We initially discretize the time interval [0, *T*] into *B* intervals of length *T*/(*B* − 1). As we generate new samples of **t**, we expand or contract our grid according to the current value of *t*_2_ by keeping the grid interval length fixed to *T*/(*B* − 1), effectively increasing or decreasing the dimension of *γ*.

#### 2.8.4 Local updates of coalescent times

In addition to HMC updates of coalescent times, we propose a move of a single coalescent time (excluding the TMRCA *t*_2_) chosen uniformly at random and sampled uniformly in the intercoalescent interval; that is, we choose *i* ~ *U*({*n, n* − 1,…, 3}) and 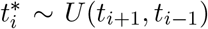. This is a symmetric proposal and the corresponding Metropolis-Hastings acceptance probability is

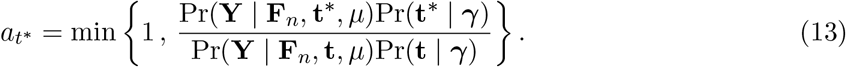

While these updates may seem unnecessary in light of the Hamiltonian updates of coalescence times (Section 2.8.3), we observed better performance of our MCMC sampler by including this additional proposal. One reason may be our choice of **M** in section 2.8.3 that does not account for the geometric structure of the posterior distribution of coalescent times. However, a better choice of M comes with higher computational burden than a simple local update of coalescent times.

#### 2.8.5 Multiple Independent loci

Thus far, we have assumed our data consist of a single linked locus of s segregating sites. We can extend our methodology to *l* independent loci with *s_i_* segregating sites for *i* = 1,…,*l*. In this case, our data 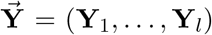 consist of *l* aligned sequences with elements {0,1}, where 0 indicates the ancestral allele as before. We then jointly estimate the Tajima’s genealogies 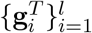, precision parameter *τ*, and vector of log effective population sizes *γ* through their posterior distribution:

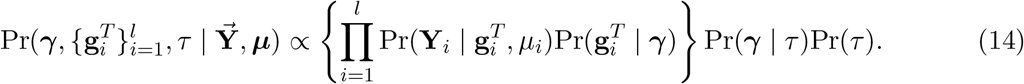

In Equation (14), we enforce that all loci follow the same effective population size trajectory but every locus can have its own mutation rate *μ_i_*.

## 3 Results

### 3.1 The performance of BESTT in applications to simulated data

We tested our new method, BESTT, on simulated data under four different demographic scenarios. Note that in this section, *N*(*t*) is rescaled to the coalescent time scale, meaning that 1/*N*(*t*) is the pairwise rate of coalescence at time *t* in the past relative to the rate at the present time zero. We simulated genealogies under four different population size trajectories:

1. A period of exponential growth followed by constant size:

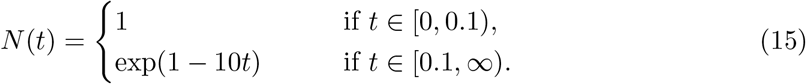
2. A trajector with instantaneous growth:

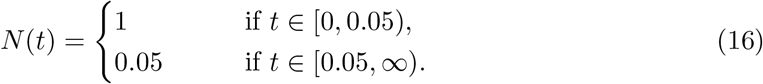
3. An exponential growth: *N*(*t*) = 25*e*^−5*t*^
4. A constant trajectory: *N*(*t*) = 1

Given a genealogy of length 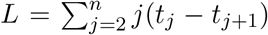, where *t_j_* − *t*_*j*+1_ is the intercoalescent length while there are *j* lineages, we drew the total number of mutations (segregating sites) *s* according to a Poisson distribution with parameter *μL*. We then placed the mutations uniformly at random along the branches of the genealogy. For each of the *s* mutations, we assigned the mutant type to individuals descending from the branch where the mutation occurred and the ancestral type otherwise.

We summarize our posterior inference 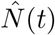 by the posterior median and 95% Bayesian credible intervals after 200 thousand iterations and thinned every 10 iterations with 100 iterations of burn in. Our initial number of change points for *N*(*t*) was set to 50 over the time interval (0,*t*_2_), where *t*_2_ is the initialized time to the most recent common ancestor; however, over the course of MCMC iterations, this number could increase or decrease according to the posterior distribution of *t*_2_.

We assess accuracy and precision of our estimates using the sum of relative errors (SRE)

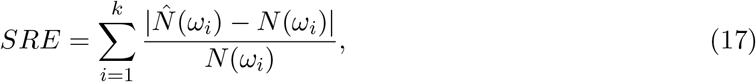

where 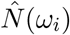 is the estimated effective population size trajectory at time *ω_i_*. Second, we computed the mean relative width as

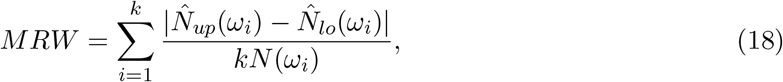

where 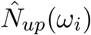 corresponds to the 97.5% upper limit and 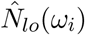 corresponds to the 2.5% lower limit of the estimated posterior distribution of *N*(*ω_i_*). In addition, we measured how well the 95% credible intervals cover the truth and compute the envelope measure, ENV:

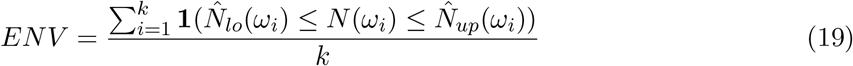

We first simulated 3 datasets of *n* = 10 individuals with an average number of 100 segregating sites under different types of population size trajectories: constant, exponential growth and instantaneous growth. Results are depicted in the first column of Figure 8. Posterior medians and 95% credible intervals are shown as black curves and gray shaded areas respectively. The trajectory used to simulate the data is depicted as a dashed line. Figure 8 shows that our BESTT method recovers the constant and exponential growth trajectories very well but the instantaneous growth scenario is less accurate and with high uncertainty (wide credible intervals). In all three cases, our envelope measure is above 95%. Performance measures on all simulations are summarized in Table 1.

**Table 1:** Empirical measures of performance in the simulations described in the text

| Simulation | % ENV | SRE | MRW |
| --- | --- | --- | --- |
| Instantaneous growth($n=10,s=31$) | 96 | 5.87 | 124352 |
| Instantaneous growth ($n=10,s=63$) | 100 | 2.15 | 2296 |
| Instantaneous growth ($n=10,s=120$) | 98 | 0.53 | 80 |
| Instantaneous growth ($n=25$) | 90 | 0.40 | 3.43 |
| Instantaneous growth ($n=35$) | 92 | 0.31 | 3.16 |
| Constant | 100 | 0.30 | 1.16 |
| Exponential | 100 | 0.35 | 5.45 |
| Exp. & const. ($n=10, 1$ locus) | 100 | 4.31 | 22608 |
| Exp. & const. ($n=10, 5$ loci) | 100 | 2.37 | 309.1 |
| Exp. & const. ($n=10, 10$ loci) | 100 | 0.16 | 4.19 |

We analyzed the effect of increasing the number of segregating sites, the number of samples and the number of independent genealogies on posterior inference with BESTT. In all three cases, we expect our method to better recover the truth. Figure 5 shows our results on simulated data under a population size trajectory with instantaneous growth (Equation 16) of *n* = 10 individuals with 31, 63 and 120 segregating sites. As expected, our method recovers the truth with higher precision (MRW) and accuracy (SRE) when we increase the number of segregating sites. Increasing the number of segregating sites may result in more constraints in the gene tree. For *n* = 10, there are 7936 possible ranked tree shapes, however for the datasets simulated with 31, 63 and 102 segregating sites, there are only 2582 ± 32, 2670 ± 34 and 556 ± 7 ranked tree shapes compatible with their corresponding gene trees. These numbers were estimated by importance sampling (Cappello and Palacios, 2019).

**Figure 5:**
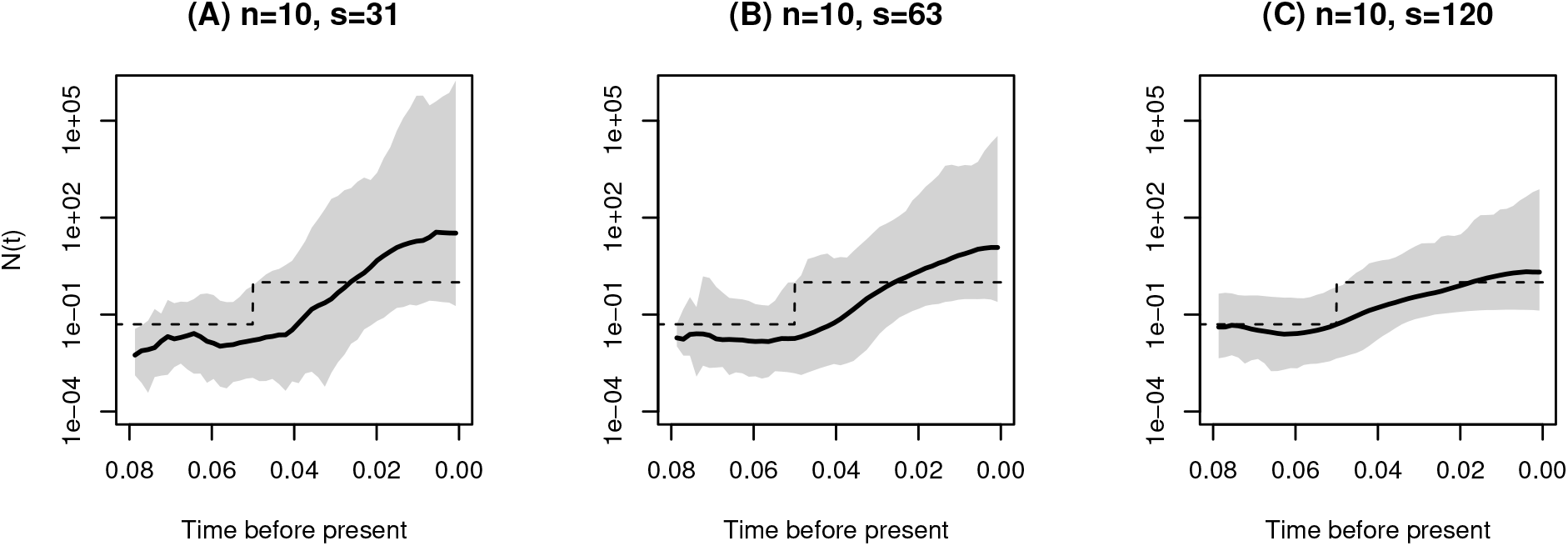
Varying the number of segregating sites. Posterior inference from simulated data of *n* = 10 sequences under a population size trajectory with instantaneous growth (dashed lines). s is the number of segregating sites. Posterior medians are depicted as solid black lines and 95% Bayesian credible intervals are depicted by shaded areas.

As another performance assessment, we simulated datasets from a population size trajectory with instantaneous growth with varying number of samples. We simulated datasets with *n* = 10, 25 and 35 samples with 215 expected number of segregating sites. Our results depicted in Figure 6 show that our method performs better in terms of SRE and MRE when the number of samples increases. Similarly, precision (MRW) and accuracy (SRE) increases when inference is done from a larger number of independent datasets. Finally, Figure 7 shows our results from 1, 5 and 10 datasets simulated from 1, 5 and 10 independent genealogies of 10 individuals with a population size trajectory of growth followed by a constant period (Equation 15). As expected, our method’s performance substantially increases by increasing the number of independent datasets.

**Figure 6:**
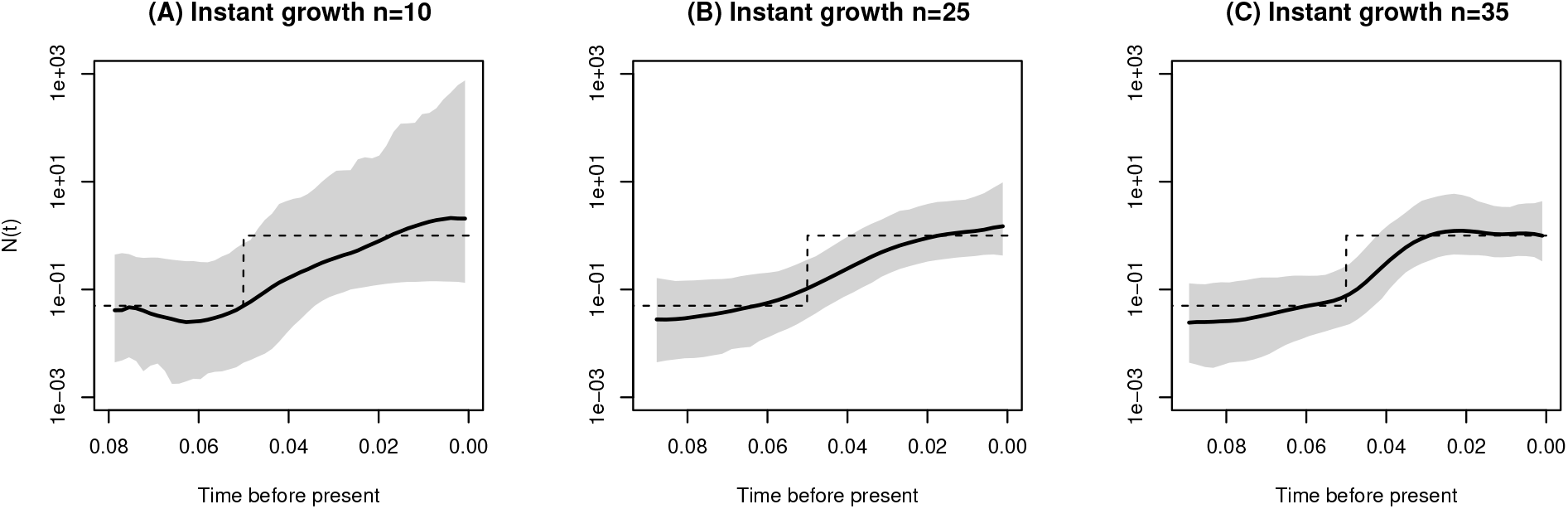
Varying the number of samples under a population size trajectory with instantaneous growth. Posterior inference from simulated data of *n* = 10, 25 and 35 sequences under the population size trajectory with instantaneous growth. Shaded areas correspond to 95% credible intervals, solid lines to posterior median and dashed line to the truth.

**Figure 7:**
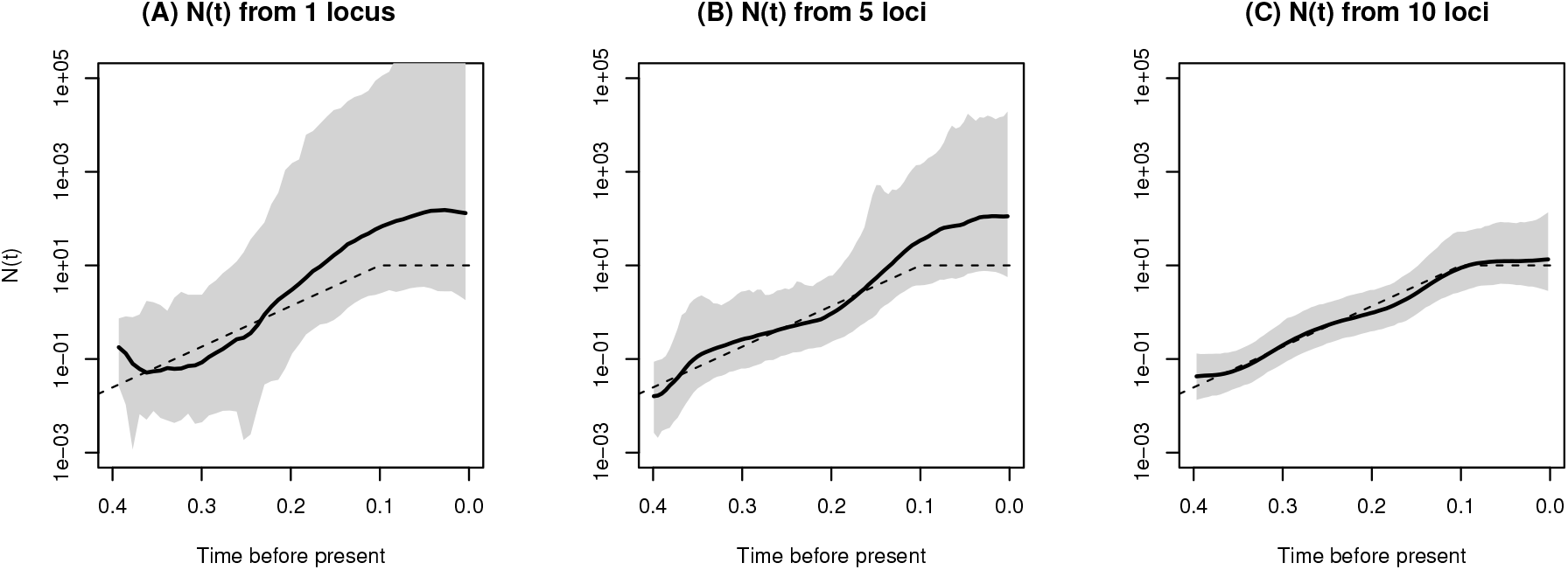
Multiple independent datasets. Posterior inference from simulated data of *n* = 10 sequences under exponential followed by constant trajectory (eq. 15). (**A**) Inference from a single simulated dataset, (**B**) from 5 independently simulated datasets, and (**C**) from 10 independently simulated datasets. Shaded areas correspond to 95% credible intervals, solid black lines show posterior medians and dashed lines show the simulated truth.

### 3.2 Comparison to other methods

To our knowledge, there is no other method for inferring (variable) effective population size over time from haplotype data that assumes the infinite sites mutation and a nonparametric prior on *N*(*t*), therefore we cannot have a direct comparison of our method to others. Moreover, our method is the only one that explicitly averages over Tajima genealogies instead of Kingman genealogies. BEAST (Drummond et al., 2012) is a program for analyzing molecular sequences that uses MCMC to average over the Kingman tree space and it is therefore a good reference for comparison to our method. We compared our results to the Extended Bayesian Skyline Plot method (EBSP) (Heled and Drummond, 2008) and the Skygrid method (Gill et al., 2013) implemented in BEAST.

Since the infinite sites mutation model is not implemented in BEAST, we first converted our simulated sequences of 0s and 1s to sequences of nucleotides by sampling *s* ancestral nucleotides uniformly on {*A, T, C, G*} and assigning one of the remaining 3 types uniformly at random to be the mutant type. This corresponds to a simulation of the Jukes-Cantor mutation model (Jukes and Cantor, 1969) that is currently implemented in BEAST.

We compare the results of BESTT to those of BEAST EBSP and Skygrid (Drummond et al., 2005, 2012) in Figure 8. We note that results from BEAST are generated from 10 million iterations and thinned every 1000 iterations, while results from BESTT are generated from 200 thousand iterations.

**Figure 8:**
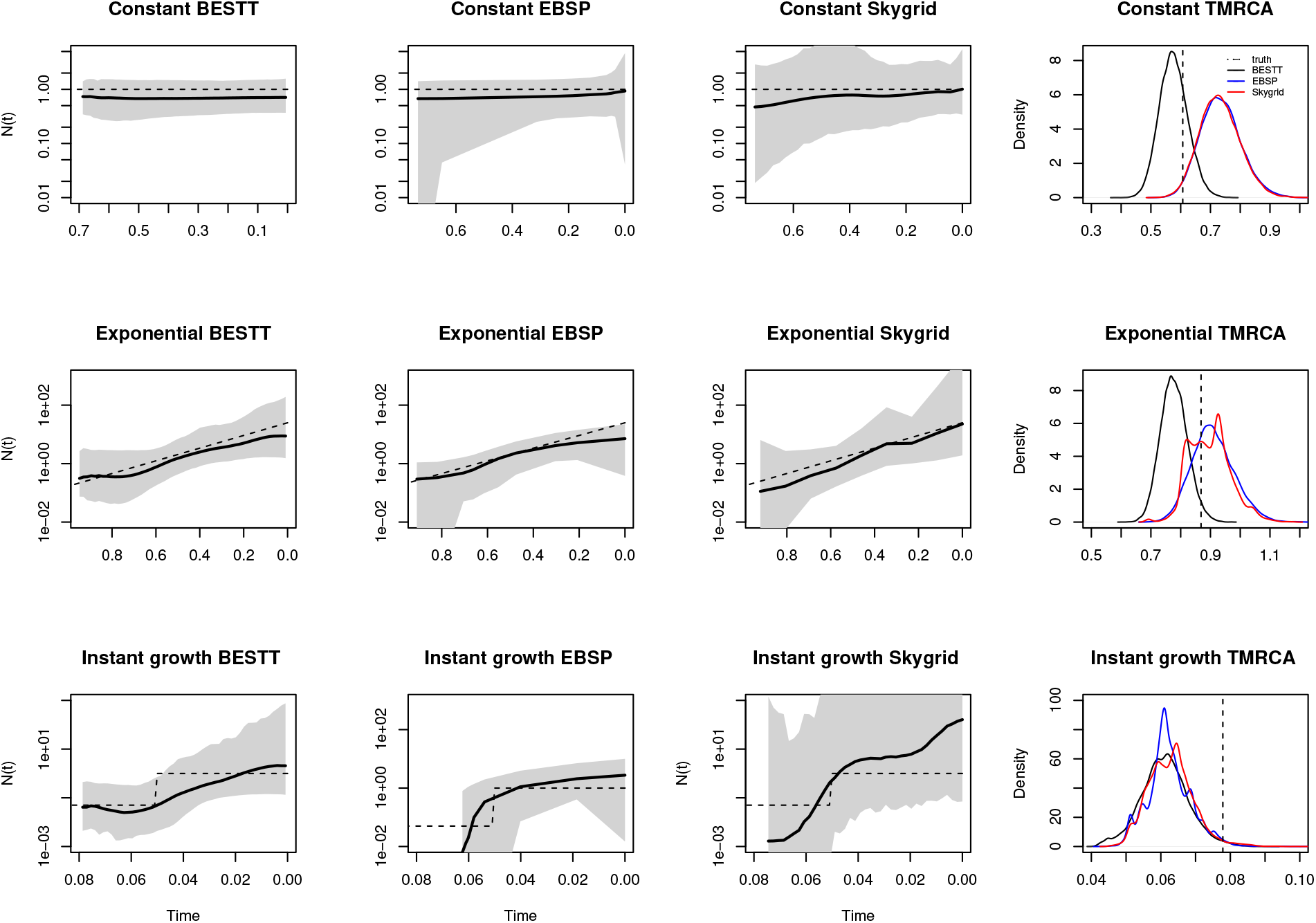
BESTT and BEAST Comparison. Posterior inference from simulated data of *n* = 10 sequences under constant, exponential and instantaneous growth trajectories (rows) obtained from our method BESTT (first column), BEAST EBSP (second column) and BEAST Skygrid (third column). Shaded areas correspond to 95% credible intervals, solid black lines show posterior medians and dashed lines show the simulated truth. In the fourth column, we show the posterior density of the time to the most recent common ancestor (TMRCA) from the three methods: BESTT (black), BEAST EBSP (blue) and BEAST Skygrid (red). The true value of the TMRCA is depicted as a vertical dashed line.

We compared our point estimates 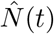 from all methods to the ground truth for each simulation (Table 2). In two cases, BESTT has better envelope than BEAST. For the exponential growth simulation (Figure 8, second row) the BEAST EBSP result has better SRE and MRW, however, the credible intervals are uneven with very wide intervals at the ends. In all cases, the BEAST Skygrid results have wider credible intervals. For the instantaneous growth simulation (Figure 8, third row), BEAST EBSP did not generate many simulations beyond the time point 0.06, for this reason we recomputed the performance statistics for the overlapping time interval (0, 0.06). In this interval, BESTT outperforms both methods implemented in BEAST in terms of envelope and SRE. The last column of Figure 8 shows the posterior distribution of the time to the most recent common ancestor (TMRCA). For the case of constant population size, the true value of the TMRCA is contained in the 95% BCI estimated with BESTT but it is not contained in the 95% BCIs estimated with the two methods implemented in BEAST. In the exponential growth simulation, the true TMRCA is contained in the 95% BCIs estimated with the three methods and the instant growth method, the true TMRCA is not contained in the 95% BCIs of the three methods.

**Table 2:** Performance comparison between BESTT and BEAST in simulations

| Simulation | % ENV |  |  | SRE |  |  | MRW |  |  |
| --- | --- | --- | --- | --- | --- | --- | --- | --- | --- |
|  | BESTT | EBSP | Skygrid | BESTT | EBSP | Skygrid | BESTT | EBSP | Skygrid |
| Constant | **100** | **100** | **100** | 0.3 | **0.24** | **0.24** | **1.16** | 1.49 | 4.41 |
| Exponential | **100** | 97 | **100** | 0.35 | **0.26** | 0.29 | 5.45 | **2.56** | 46.13 |
| Inst. growth | 97 | 94 | **100** | **0.61** | 2.65 | 22.6 | 105.6 | **14.95** | >1000 |

We note that BEAST Bayesian Skygrid (Gill et al., 2013) is a more comparable method to BESTT since it assumes Gaussian process priors on log *N*(*t*) like BESTT.

### 3.3 Computational performance of BESTT

BESTT approximates the posterior distribution (a) Pr((*N*(*t*))_*t*≥0_,**g**^*T*^, ***τ*** | **Y**, *μ*), where **g**^*T*^ is a Tajima’s genealogy instead of (b) Pr((*N*(*t*))_*t*≥0_, **g, *τ*** | **Y**\*, *μ*), where **g** is a Kingman’s genealogy and **Y**\* is the labeled data, in order to estimate (*N*(*t*))_*t*>0_. These two posterior distributions are the same when every individual of the sample has its own private mutation group and no shared mutation groups. Otherwise, the number of Tajima’s trees compatible with observed data **Y**, *i.e*. Tajima’s trees **g**^*T*^ such that Pr(**g**^*T*^ | **Y**) > 0, is smaller than the number of Kingman’s trees compatible with observed labeled data **Y**\* (Cappello and Palacios, 2019). That is, we are required to estimate the posterior of a smaller number of trees. For this reason, we argue that Tajima’s coalescent is a more efficient model than Kingman’s coalescent for estimating the posterior distribution of (*N*(*t*))_*t*≥0_. However, a single conditional likelihood calculation Pr(**Y** | **g**^*T*^, *μ*) requires the sum over all possible allocation of mutation groups to branches of **g**^*T*^. Our algorithm only accounts for allocations constrained by the DAG and the ranked tree shape of **g**^*T*^. For the data depicted in Figure 2A,B and **g**^*T*^ of Figure 2C, there are only 8 different possible allocation “paths” of all mutation groups to branches. In Appendix C we detail how our algorithm finds these paths. The number of paths depends on the number of subtrees with the same family size path in the DAG and in the ranked tree shape. In the best case, our algorithm will find a path in *O*(*no*), where no is the number of nodes in the gene tree. In general, the number of allocation paths will be much smaller than the number of labeled trees compatible with a ranked tree shape. In our implementation, we estimate posterior (a) with MCMC. The main difference between our MCMC algorithm and the MCMC algorithm implemented in BEAST is the tree topology sampler. While our MCMC algorithm explores the space of ranked tree shapes with local move proposals of ranked tree shapes, BEAST explores the space of labeled, ranked tree shapes with local move proposals of labeled trees. A formal assessment of the efficiency of our MCMC algorithm and its comparison to the MCMC implementation in BEAST is beyond the scope of this manuscript and subject of future research.

## 4 Inferring human population demography from mtDNA

We selected *n* = 35 samples of mtDNA at random from 107 Yoruban individuals available from the 1000 Genomes Project phase 3 (The 1000 Genomes Project Consortium, 2015). We retained the coding region: base pairs 576 – 16, 024 according to the rCRS reference of Human Mitochondrial DNA (Anderson et al., 1981; Andrews et al., 1999) and removed 38 indels. Of the 260 polymorphic sites, we retained 240 sites compatible with the infinite sites mutation model. The final file is available in https://github.com/JuliaPalacios/phylodyn. To encode our data as 0s and 1s, we use the inferred root sequence **RSRS** of Behar et al. (2012) to define the ancestral type at each site. To rescale our results in units of years, we assumed a mutation rate per site per year of 1.3 × 10^−8^ (Rebolledo-Jaramillo et al., 2014). We compare our results with the Extended Bayesian Skyline method (Drummond et al., 2012) implemented in BEAST in Figure 9. When applying BEAST, we assumed the Jukes-Cantor mutation model. Both methods detect an inflection point around 20kya followed by exponential growth. The mean time to the most recent ancestor (TM-RCA) inferred for these YRI mtDNA samples with BESTT is around 170kya with a 95% BCI of (142868,207455), while the mean TMRCA inferred with BEAST is around 160kya with a 95% BCI of (133239,196900). In Appendix D, we include two more comparisons of BESTT and BEAST.

**Figure 9:**
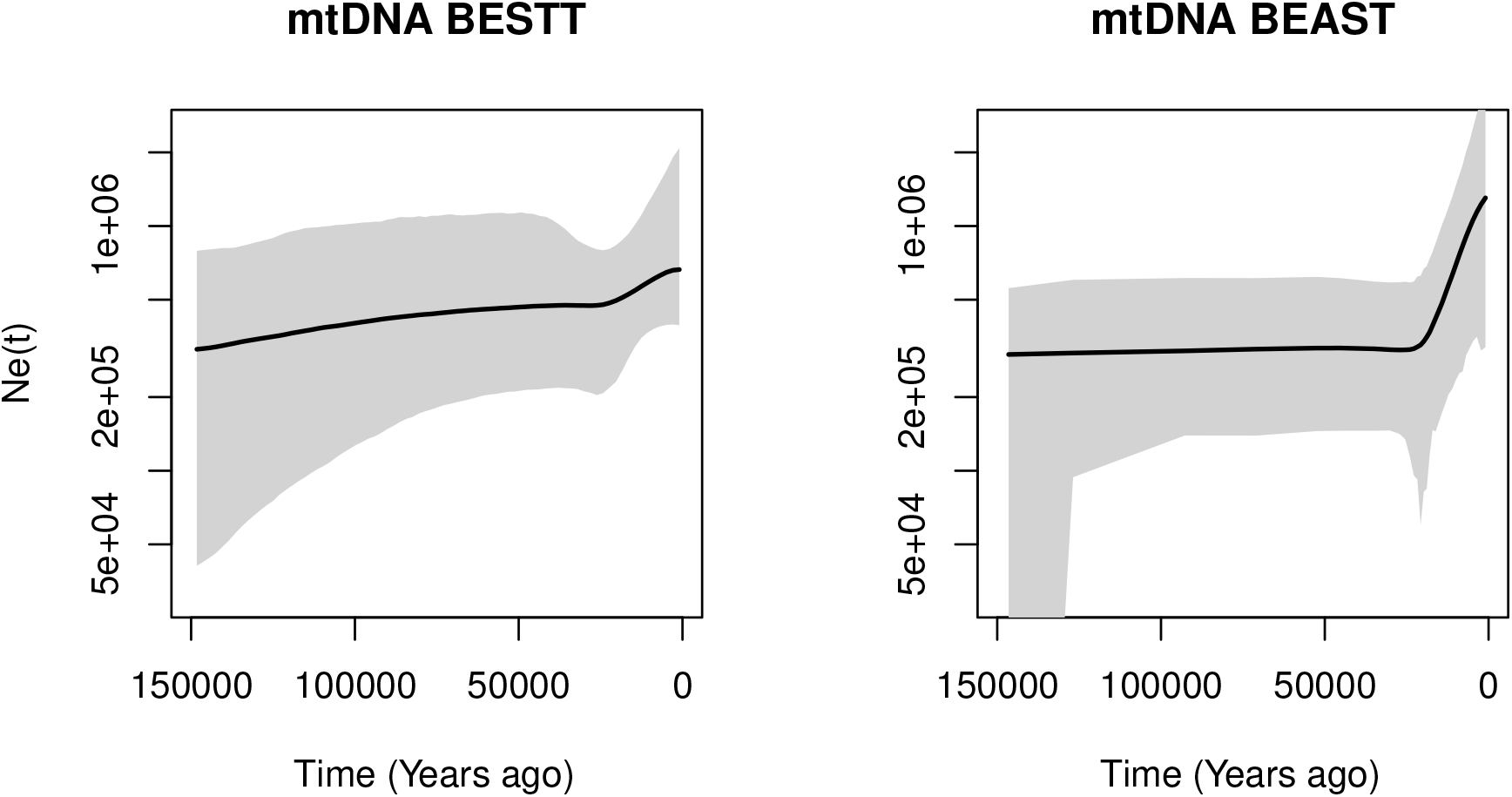
Posterior inference of female effective population size from 35 mtDNA samples from Yoruban individuals in the 1000 Genomes Project using our method BESTT (first plot) and the BEAST Extended Bayesian Skyline Plot (second plot). Posterior median curves are depicted as solid black lines and 95% credible intervals by shaded regions.

## 5 Discussion

The size of emergent sequencing datasets prohibits the use of standard coalescent modeling for inferring evolutionary parameters. The main computational bottleneck of coalescent-based inference of evolutionary histories lies in the large cardinality of the hidden state space of genealogies. In the standard Kingman coalescent, a genealogy is a random labeled bifurcating tree that models the set of ancestral relationships of the samples. The genealogy accounts for the correlated structure induced by the shared past history of organisms and explicit modeling of genealogies is fundamental for learning about the past history of organisms. However, the genomic era is producing large datasets that require more efficient approaches that efficiently integrate over the hidden state space of genealogies.

In this manuscript we show that a lower resolution coalescent model on genealogies, the “Tajima’s coalescent”, can be used as an alternative to the standard Kingman coalescent model. In particular, we show that the Tajima coalescent model provides a feasible alternative that integrates over a smaller state space than the standard Kingman model. The main advantage in Tajima’s coalescent is to model the ranked tree topology as opposed to the fully labeled tree topology as in Kingman’s coalescent.

*A priori*, the cardinality of the state space of ranked tree shapes is much smaller than the cardinality of the state space of labeled trees. However, in this manuscript we show that when the Tajima coalescent model is coupled with the infinite sites mutation model, the space of ranked tree shapes is constrained by the data and the reduction on the cardinality of the hidden state space of Tajima’s trees is even more pronounced than expected.

In order to leverage the constraints imposed by the data and the infinite-sites mutation model, we apply Dan Gusfield’s perfect phylogeny algorithm (Gusfield, 1991) to represent sequence alignments as a gene tree. We exploit the gene tree representation for conditional likelihood calculations and for exploring the state space of ranked tree shapes.

For the calculation of the likelihood of the data conditioned on a given Tajima’s genealogy, we augment the gene tree representation of the data with the Tajima’s genealogy and map observed mutations to branches. We define a directed acyclic graph (DAG) with the augmented gene tree. This new representation as a DAG allows for calculating the likelihood as a backtracking algorithm that transverses the gene tree from the leaves to the root. Our implementation’s computational bottleneck lies in the likelihood calculation. Given a Tajima’s genealogy, our likelihood algorithm sums over all possible allocation of mutation groups to branches. Although this number is generally much smaller than the number of labeled genealogies, our algorithm can be further optimized. In future studies, we will explore as sum-product type of algorithm for the likelihood calculation. In the present implementation we are able to infer effective population size trajectories from samples of size *n* ≈ 35 in a regular personal laptop computer within few hours.

Our statistical framework draws on Bayesian nonparametrics. We place a flexible geometric random walk process prior on the effective population size that allows us to recover population size trajectories with abrupt changes in simulations. The inference procedure proposed in this manuscript relies on Markov chain Monte Carlo (MCMC) methods with 3 large Gibbs block updates of: coalescent times, effective population size trajectory and ranked tree shape topology. We use Hamiltonian Monte Carlo updates for continuous random variables: coalescent times and effective population size; and a Metropolis Hastings sampler for exploring the space of ranked tree shapes. For exploring the genealogical space, Markovtsova et al. (2000) suggest a joint local proposal for both coalescent times and topology. Here we restrict our attention to the topology alone. A future line of research includes the development of a joint local proposal of coalescent times and ranked tree shapes. We also envision that a joint sampler of coalescent times and effective population size trajectories should improve mixing and convergence.

Our method does not model recombination, population structure or selection. It assumes completely linked and neutral segments from individuals from a single population, and the infinite sites mutation model. While this model is a good approximation for some human molecular data, it is not appropriate for modeling molecular data from other organisms such as pathogens and viral populations. Finally, haplotype data of many organisms is usually sparse with few unique haplotypes presented at high frequencies. Since our algorithm exploits molecular data at the haplotype level, our proposed method is ideally suited for this scenario where the space of ranked tree shapes is drastically smaller than the space of labeled topolgies.

## Acknowledgements

The authors thank the editor and two anonymous referees whose suggestions considerably improved the manuscript. This research is supported in part by a National Institutes of Health grant R01-GM-131404 and the Alfred P. Sloan Foundation to J.A.P.. We want to acknowledge the developers of `R-ape, R-phangorn` and `R-phylodyn` that facilitated our implementations. A.V. was supported in part by the chaire program Modélisation Mathématique et Biodiversité of Veolia Environnement - École Polytechnique - Museum National d’Histoire Naturelle - Fondation X. A.V. and J.A.P. was supported by the France-Stanford Center for interdisciplinary Studies. This was work also supported by the National Science Foundation CAREER Award DBI-1452622 to S.R.

## Appendix A Matrix representation of a ranked tree shape

Our algorithms exploit the following encoding of a ranked tree shape by a triangular matrix of size *n* × *n*, which we denote by **F** (Figure 10). Recall that, by convention, *t*_*n*+1_ = 0 and *t*_1_ = +∞.

**Figure 10:**
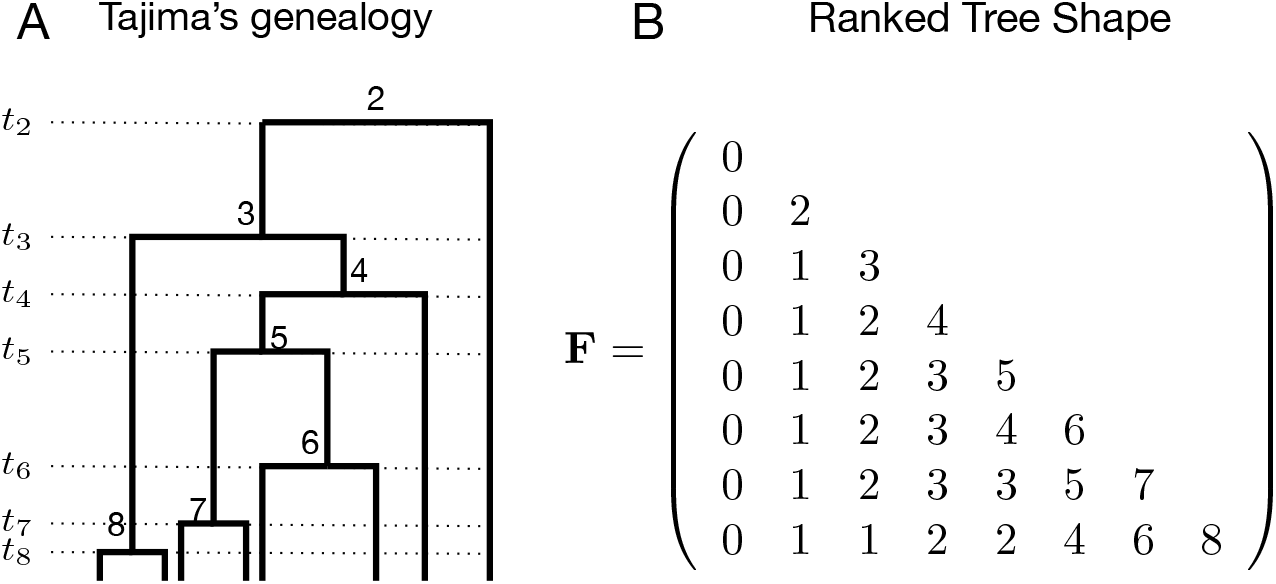
Ranked tree shape. Left: Example of a Tajima’s genealogy (redrawn from Figure 1A) with coalescent events ranked from 2 at time *t*_2_ to *n* at time *t_n_*. Right: The corresponding **F**_*n*_ matrix, with *n* = 8, that encodes the ranked tree shape information of the Tajima’s genealogy on the left. **F**_*i,j*_ denotes the number of lineages that do not coalesce in the time interval (*t*_*i*+1_, *t_j_*). In particular, **F**_*n,i*_ for *i* in {2, 3,…,*n*} denotes the number of singletons (external branches that have not coalesced) in the time interval (*t*_*i*+1_, *t_i_*).

First, we declare that **F**_*i,j*_ = 0 if *j* > *i*. Next, the number of lineages through time is encoded on the diagonal of **F**: **F**_*i,i*_ = *i* for *i* in {2, 3,…, *n*}. Finally, for *j* < *i*, the entry **F**_*i,j*_ denotes the number of lineages that do not coalesce in the time interval (*t*_*i*+1_, *t_j_*); in particular, **F**_*i*,1_ = 0 and for every *i* in {2,3,…, *n*}, **F**_*n,i*_ denotes the number of singletons (*i.e*., external branches that have not coalesced) in the time interval (*t*_*i*+1_, *t_i_*) (Figure 10). Other statistics of the ranked tree shape can be expressed in terms of the corresponding matrix **F**. Among them, the number *c* of cherries is equal to the number of times that the number of singletons decreases by 2 between lines *i* and *i* − 1, since such an event means that the coalescence separating these two epochs was that of two external branches. That is,

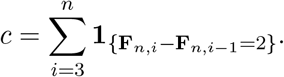

## Appendix B Detailed allocation of mutation groups along **g**^*T*^

The latent allocation random variables {*A_j_*} are constrained by the information in the Tajima’s genealogy **g**^*T*^. In a given **g**^*T*^, every subtree is labeled by its ranking from past to present (Figure 10). Subtree *i* is subtended by branch *b_i_* with length *l_i_*, for *i* = 2,…, *n*. We will assume that *l*_2_, the length of the root branch, is 0. Let *c* be the number of cherries (nodes with two leaves) in **g**^*T*^; the two branches of a given cherry share the same label *b_j_* ∈ {*b*_*n*+1_,…,*b*_*n*+*c*_}. The actual label of external branches is arbitrary but, for ease of exposition in our figures, we first label the cherries’ branches from left to right by {*b*_*n*+1_,…, *b*_*n*+*c*_}; singleton branches are labeled from left to right by *b*_*n*+*c*+1_,…, *b*_2*n*−*c*_ (Figure 2C). As mentioned before, the length of *X_j_* is the number of the corresponding sister nodes in 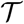 that were grouped together in forming node *Z_j_*. In this case, *A_j_* = (*A*_*j*,1_,…, *A*_*j*,|*ch*(*j*)|_) denotes a collection of |*ch*(*j*)| vectors of branch labels in **g**^*T*^ subtending the child-node subtrees of node *Z_j_*. *A*_*j*,1_ corresponds to the branch subtending from the leftmost child node of *Z_j_* on **D**, *A*_*j*,2_ corresponds to the branch subtending from the next child node of *Z_j_*, etc., and *A*_*j*,|*ch*(*j*)|_ corresponds to the branch subtending from the rightmost child node of *Z_j_* on **D**. Observe that, since we group some of the leaf nodes in 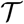 into a single node in **D**, any *A_j,k_* may be a vector of branch labels; for example *A*_1,1_ = (*b*_12_, *b*_9_) and *A*_1,2_ = *b*_10_ in Figure 3B.

## Appendix C Algorithms for conditional likelihood calculation

The following two algorithms detail the calculation of Pr(**Y**| *g^T^*, *μ*). **Y** is encoded in *GeneTree*, the observed data as a Tree structure. Each node in *GeneTree* has number of descendants (or lineages) and mutation information attached to it. Tajima’s genealogy *g^T^* is encoded as *Fpath* that contains the ranked tree shape **F***_n_* and times that contains the vector of coalescent times **t** multiplied by the mutation rate *μ*.

### Algorithm 1 Calculate Likelihood(*Fpath*, *times*, *GeneTree*) procedure

~~~
**Input**: *GeneTree*, *FPath*
**Output**: Log Likelihood *LL*
 1: Initiate *pool* to be the set of leave nodes of *GeneTree* with at least one descendant. Initiate *LL* and *index* to be zero. Initiate *current_path* to be empty.
 2: Call CalcLL_recursive(*LL*, *index*, *current_path*, *Fpath*, *times*, *Genetree*).
 3: **return** *LL*
~~~

### Algorithm 2 CalcLL_recursive(*LL*, *index*, *current_path*, *Fpath*, *times*, *Genetree*) procedure

~~~
1: **if** *index* = *len*(*path*) {When a complete path node is found} **then**
2:  **for** node in tree **do**
3:   Calculate log likelihood based on times and number of mutations of *node* in *current_path*.
4:   Accumulate to total log likelihood *LL*
5:  **end for**
6:**else**
7:  **for** *node in pool* **do**
8:   Check compatibility of the *node*, according to the given *Fpath*.
9:   **if** *node* is compatible with *Fpath* then
10:    Update *node* by assigning it to the current step in *Fpath*
11:    Update *pool*. If a *node* has been mapped entirely, remove node from *pool*, update its parent *node*, and potentially add parent node to *pool* if parent node has not been entirely assigned.
12:    Append this *node* to *current_path*
13:    Call CalcLL_recursive(*LL*, *index* + 1, *current_path*, *Fpath*, *Genetree*)
14:    Restore previous *node*, *pool* and *current_path*
15:   **end if**
16:  **end for**
17:**end if**
~~~

To illustrate our algorithms 1 and 2, we use our example of Figures 2 and 3. Algorithm 1 initiates the *pool* with nodes *Z*_3_,*Z*_4_, *Z*_6_, *Z*_8_,*Z*_10_. Then, Algorithm 2 cycles through this list. Assume the first node is *Z*_8_. This node has *d*_8_ = 2 descendants and the *ancestry* is *Z*_8_ – *Z*_5_ – *Z*_1_ – *Z*_0_ with sizes:2 – 3 – 7 – 16. On the other hand, the first coalescent event (from present to past) labeled 16 in *g^T^* (Figure 3A) has ancestry with sizes: 2 – 3 – 7 – 16. Therefore, this node is *compatible*. The node is removed from the pool, its parent node added to the pool and *Z*_8_ is assigned to the path. At this time *current_path* = *Z*_8_ and the *pool* becomes: *Z*_3_, *Z*_4_, *Z*_5_, *Z*_6_, *Z*_10_. The algorithm then cycles through this list and picks *Z*_10_. This node has size ancestry 2 – 3 – 7 – 16. On the other hand the second coalescent event labeled 15 has size ancestry: 2 – 3 – 5 – 7 – 9 – 16. Since the node size ancestry is contained in the second coalescent event’s size ancestry, this node is *compatible*. The current path becomes *current _path* = *Z*_8_ – *Z*_10_ and pool becomes *Z*_3_, *Z*_4_, *Z*_5_, *Z*_6_, *Z*_7_. We continue this procedure until we reach the path *current_path* = *Z*_8_ – *Z*_10_ – *Z*_3_ – *Z*_6_ – *Z*_4_ – *Z*_7_ – *Z*_5_ – *Z*_4_ – *Z*_6_ – *Z*_1_ – *Z*_2_ – *Z*_1_ – *Z*_2_ – *Z*_0_.

Once a path is found, the algorithm back tracks the path until there is one compatible node and the path continues to grow. A sequence of back tracking and growing is the following:

1. *Z*_8_ – *Z*_10_ – *Z*_3_ – *Z*_6_ – *Z*_4_ – *Z*_7_ – *Z*_5_ – *Z*_4_ – *Z*_6_ – *Z*_1_ – *Z*_2_ – *Z*_1_ – *Z*_2_ – *Z*_0_
2. *Z*_8_ – *Z*_10_ – *Z*_3_ – *Z*_6_ – *Z*_4_ – *Z*_7_ – *Z*_5_ – *Z*_4_ – *Z*_6_ – *Z*_1_ – *Z*_2_ – *Z*_1_ – *Z*_2_
3. *Z*_8_ – *Z*_10_ – *Z*_3_ – *Z*_6_ – *Z*_4_ – *Z*_7_ – *Z*_5_ – *Z*_4_ – *Z*_6_ – *Z*_1_ – *Z*_2_ – *Z*_1_
4. *Z*_8_ – *Z*_10_ – *Z*_3_ – *Z*_6_ – *Z*_4_ – *Z*_7_ – *Z*_5_ – *Z*_4_ – *Z*_6_ – *Z*_1_ – *Z*_2_
5. *Z*_8_ – *Z*_10_ – *Z*_3_ – *Z*_6_ – *Z*_4_ – *Z*_7_ – *Z*_5_ – *Z*_4_ – *Z*_6_ – *Z*_1_
6. *Z*_8_ – *Z*_10_ – *Z*_3_ – *Z*_6_ – *Z*_4_ – *Z*_7_ – *Z*_5_ – *Z*_4_ – *Z*_6_
7. *Z*_8_ – *Z*_10_ – *Z*_3_ – *Z*_6_ – *Z*_4_ – *Z*_7_ – *Z*_5_ – *Z*_4_
8. *Z*_8_ – *Z*_10_ – *Z*_3_ – *Z*_6_ – *Z*_4_ – *Z*_7_ – *Z*_5_
9. *Z*_8_ – *Z*_10_ – *Z*_3_ – *Z*_6_ – *Z*_4_ – *Z*_7_ – *Z*_5_ – *Z*_6_
10. *Z*_8_ – *Z*_10_ – *Z*_3_ – *Z*_6_ – *Z*_4_ – *Z*_7_ – *Z*_5_ – *Z*_6_ – *Z*_4_
11. *Z*_8_ – *Z*_10_ – *Z*_3_ – *Z*_6_ – *Z*_4_ – *Z*_7_ – *Z*_5_ – *Z*_6_
12. *Z*_8_ – *Z*_10_ – *Z*_3_ – *Z*_6_ – *Z*_4_ – *Z*_7_ – *Z*_5_
13. *Z*_8_ – *Z*_10_ – *Z*_3_ – *Z*_6_ – *Z*_4_ – *Z*_7_ – *Z*_5_ – *Z*_4_
14. *Z*_8_ – *Z*_10_ – *Z*_3_ – *Z*_6_ – *Z*_4_ – *Z*_7_ – *Z*_5_ – *Z*_4_ – *Z*_6_
15. *Z*_8_ – *Z*_10_ – *Z*_3_ – *Z*_6_ – *Z*_4_ – *Z*_7_ – *Z*_5_ – *Z*_4_ – *Z*_6_ – *Z*_1_
16. *Z*_8_ – *Z*_10_ – *Z*_3_ – *Z*_6_ – *Z*_4_ – *Z*_7_ – *Z*_5_ – *Z*_4_ – *Z*_6_ – *Z*_1_ – *Z*_2_
17. *Z*_8_ – *Z*_10_ – *Z*_3_ – *Z*_6_ – *Z*_4_ – *Z*_7_ – *Z*_5_ – *Z*_4_ – *Z*_6_ – *Z*_1_ – *Z*_2_ – *Z*_1_
18. *Z*_8_ – *Z*_10_ – *Z*_3_ – *Z*_6_ – *Z*_4_ – *Z*_7_ – *Z*_5_ – *Z*_4_ – *Z*_6_ – *Z*_1_ – *Z*_2_ – *Z*_1_ – *Z*_0_

The first sequence of steps 1 – 8, the path decreases. This happens because there are not alternative compatible paths until that point when the sequence starts to grow until step 10. At step 10, the algorithm does not find a compatible way to keep growing the path so the algorithm starts to back track again until step 12. From steps 12 to 18, the algorithm grows the path until a complete new path has been found. A complete path has the correspondence of coalescent events to nodes in gene tree. The first element of the path: *Z*_8_ corresponds to the coalescent event at time *t*_8_, the second element of the path *Z*_10_ corresponds to the second coalescent event at time *t*_7_. The last element of the path is *Z*_0_ when all sequences coalesce at time *t*_2_. In this example, the algorithm finds 8 paths. Once the paths are found, the algorithm computes the likelihood and the result is the sum of the likelihoods of the 8 paths.

### Markovian proposal of ranked tree shapes

The following algorithm generates a new ranked tree shape from a Markovian proposal and outputs the corresponding transition probabilities. This proposal is used in section 2.8.1.

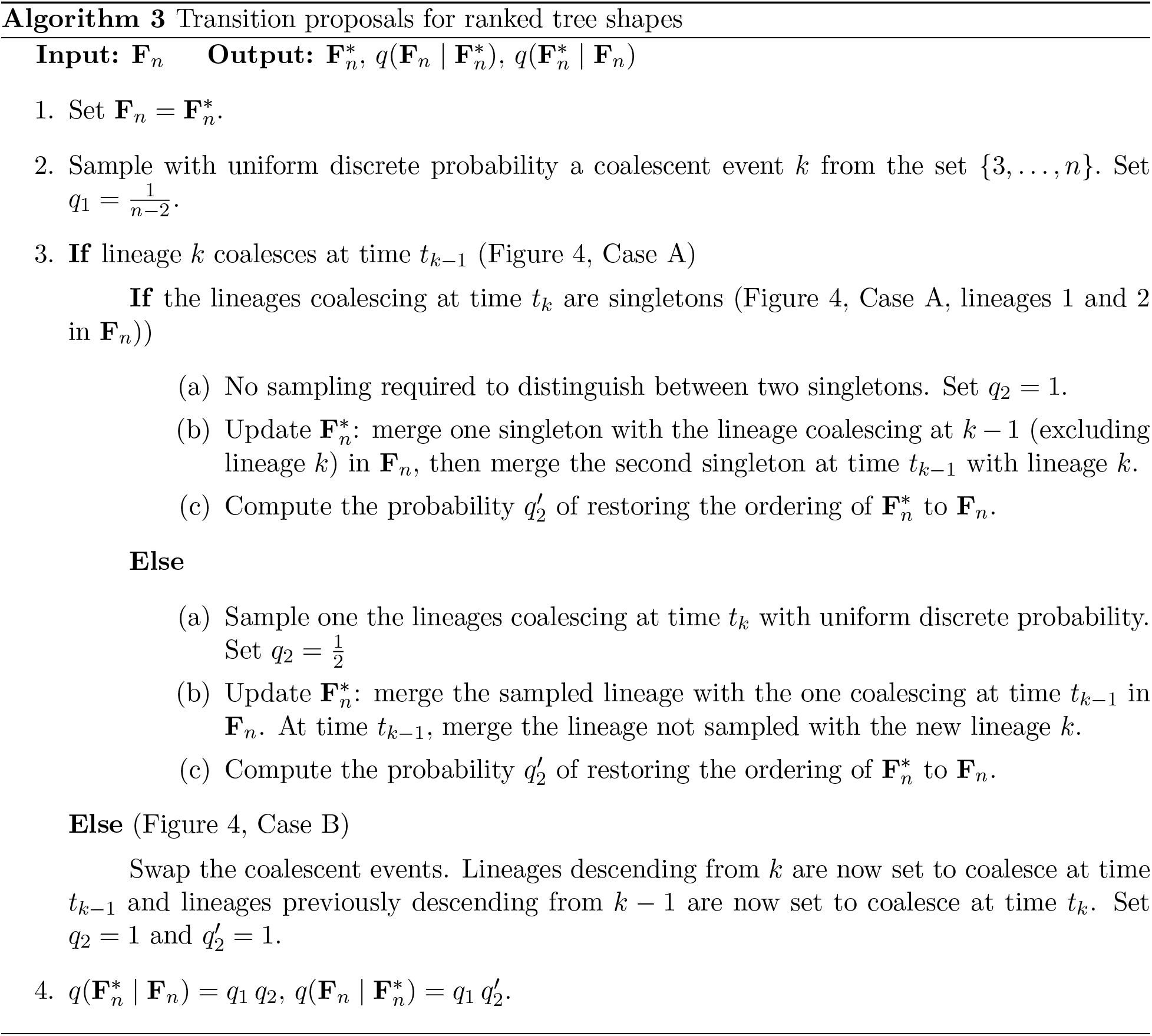

## Appendix D

We replicated the BEAST EBSP Analysis of the 35 Yoruban individuals from the 1000 Genomes Project phase 3 using the whole mtDNA coding region consisting of 15409 sites. In both cases we assumed the Jukes-Cantor mutation model (Jukes and Cantor, 1969). Figure 11 shows the comparison between EBSP inference from the 240 segregating sites retained in section 4 that are compatible with the infinite sites mutation model assumption. In both cases we recover very similar trajectories.

**Figure 11:**
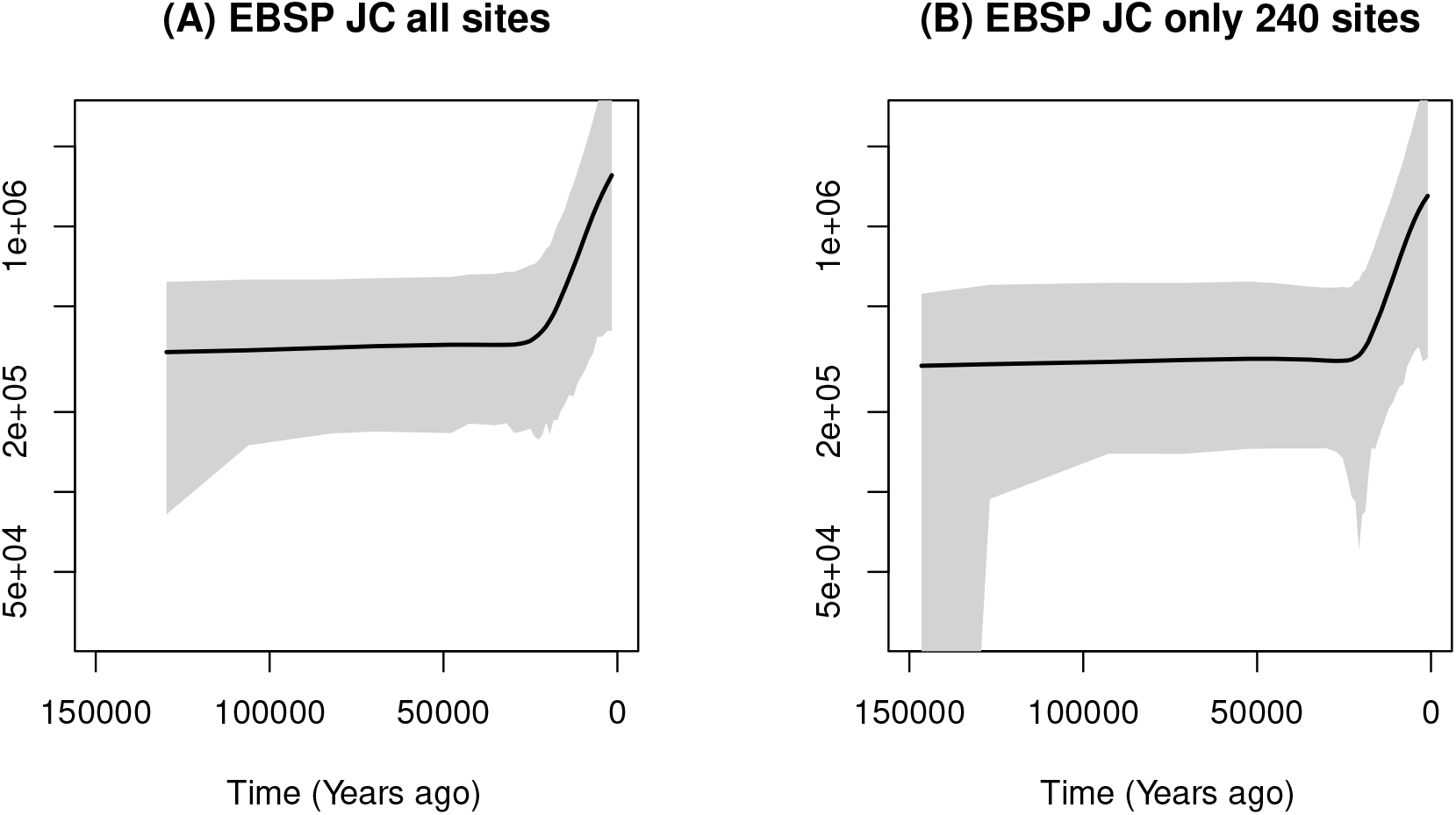
Posterior inference of female effective population size from 35 mtDNA samples from Yoruban individuals in the 1000 Genomes Project using BEAST EBSP (first plot) from all 15409 sites and the BEAST EBSP (second plot) from the 240 segregating sites retained. In both cases, the mutation model assumed is Jukes Cantor (JC). Posterior median curves are depicted as solid black lines and 95% credible intervals by shaded regions.

In addition, we compared our results with BEAST Bayesian Skyline Plot (BSP) (Drummond and Rodrigo, 2000). For our reduced dataset of 240 segregating sites, we could not generate valid inference of *N*(*t*) with Metropolis-Hastings acceptance probability greater than 0. Instead we were able to generate results with BEAST BSP from the complete dataset of 15409 sites. The comparison of our method from 240 segregating sites to BEAST BSP from 15409 sites is depicted in Figure 12.

**Figure 12:**
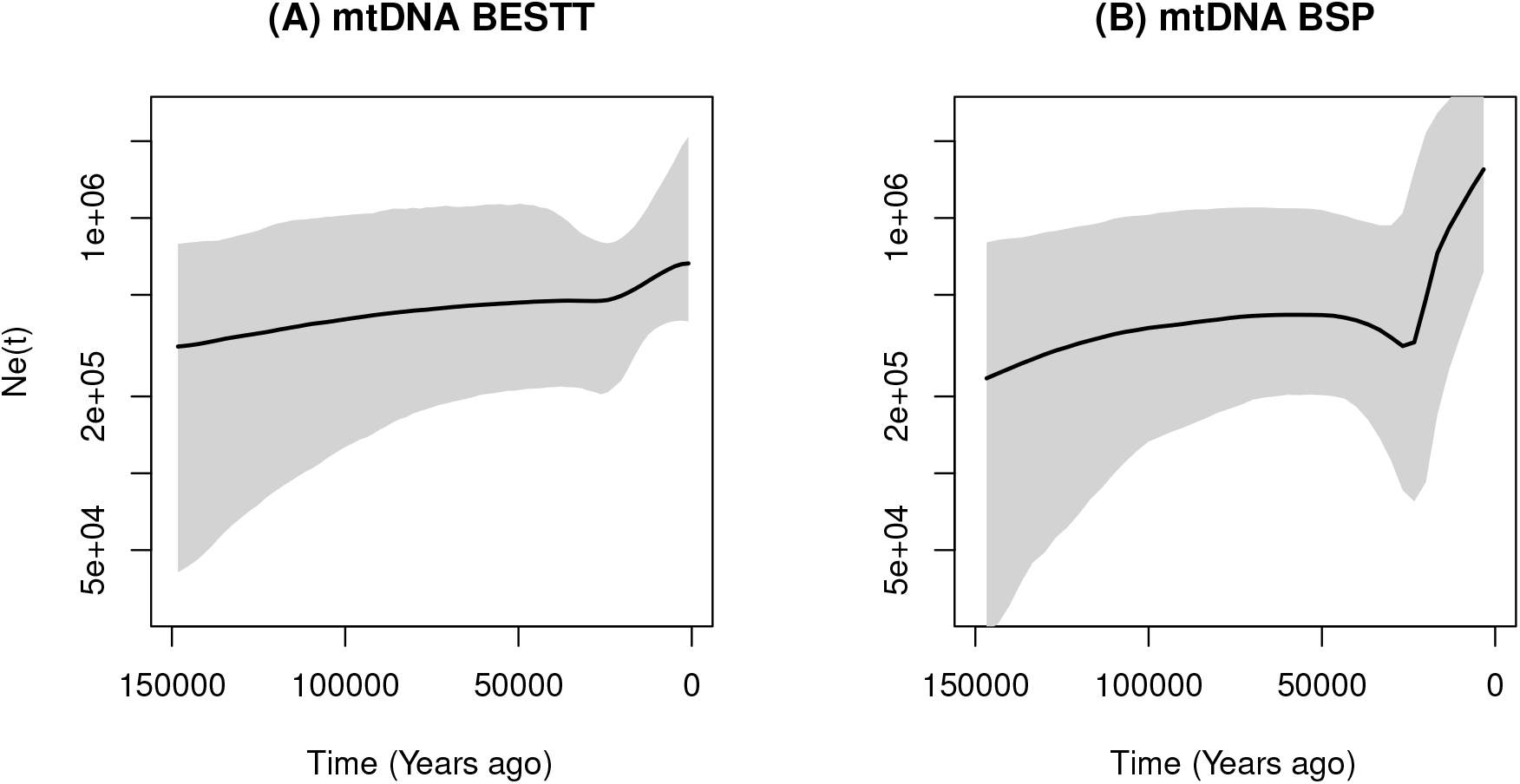
Posterior inference of female effective population size from 35 mtDNA samples from Yoruban individuals in the 1000 Genomes Project using our BESTT (first plot) from only 240 segregating sites and the BEAST BSP (second plot) from all the 15409 sites. Posterior median curves are depicted as solid black lines and 95% credible intervals by shaded regions.

## Appendix E

In Figure 13A, we show the data from Figure 2A with an additional haplotype (10) with frequency 1 and an additional column grouped with mutation group h (not shown in the table). In 13B we show the corresponding perfect phylogeny. This new perfect phylogeny has a new tip with black label 1 (frequency) subtending from a branch with 0 mutations (red label). The path from the leaf to the root shows that this haplotype has a unique mutation corresponding to mutation group *a*. We note that mutation group labels carry no information. We incorporate the labels in the Figure for ease of exposition. Since mutation group h has now multiplicity 2, the branch labeled h has now a red label 2.

**Figure 13:**
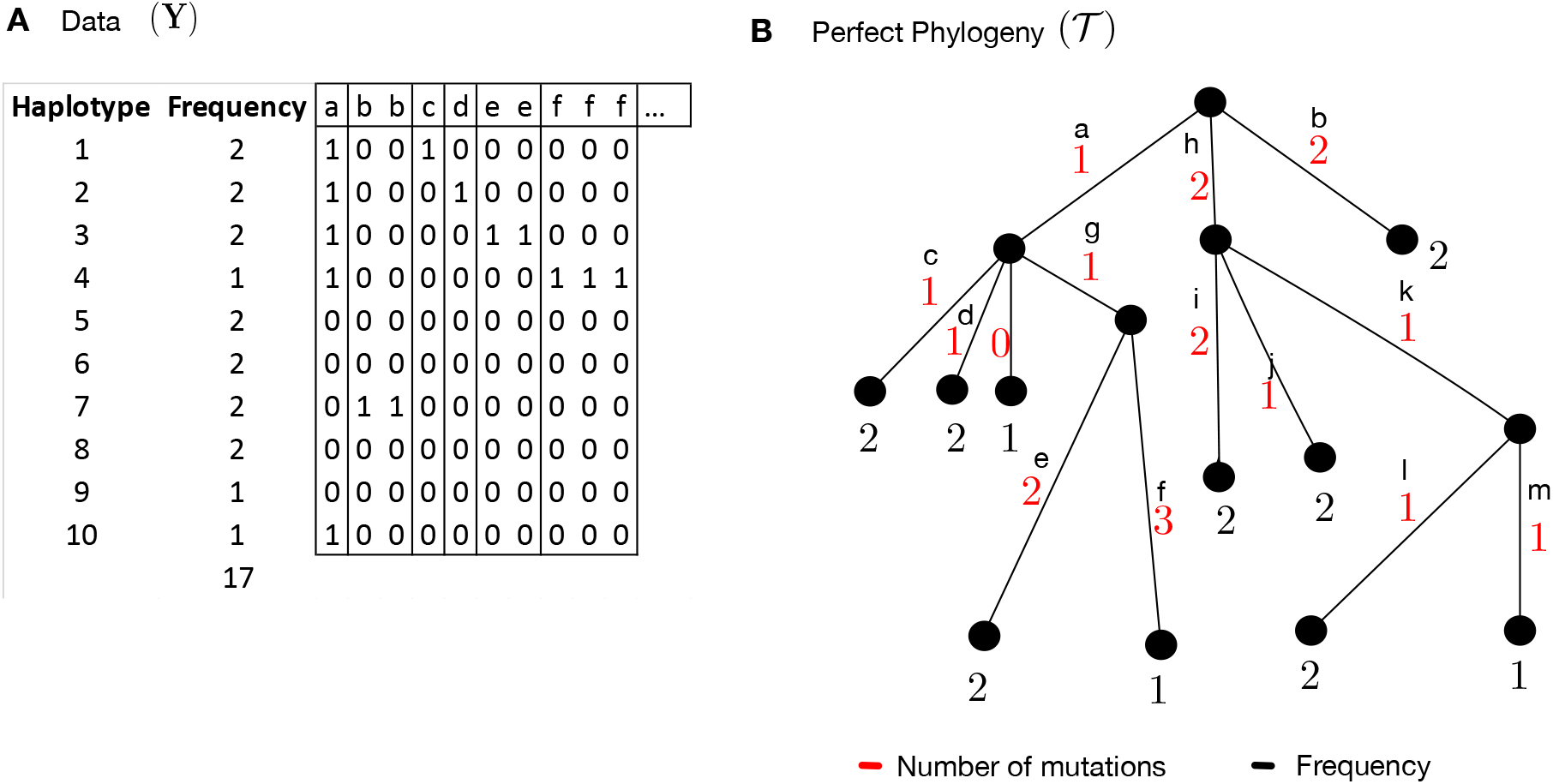
Second example of Perfect Phylogeny. **A.** Compressed data representation **Y**_*h*×*m*_ of *n* =17 sequences and *s* = 19 (columns, only the first 10 of which are shown), comprised of 10 haplotypes and 13 mutation groups. This data table has one more haplotype (10) and one more mutation labeled h than the example of Figure 2. **B.** Gene tree representation of the data in panel A. Red numbers indicate the cardinality of each mutation group (number of columns with the same label in panel A). Black letters indicate the mutation group (column labels in panel A), and black numbers indicate the frequency of the corresponding haplotype.

